# Regulatory imbalance between LRRK2 kinase, PPM1H phosphatase, and ARF6 GTPase disrupts the axonal transport of autophagosomes

**DOI:** 10.1101/2022.11.14.516471

**Authors:** Dan Dou, Erin M. Smith, Chantell S. Evans, C. Alexander Boecker, Erika L.F. Holzbaur

## Abstract

Gain-of-function mutations in the *LRRK2* gene cause Parkinson’s disease (PD), increasing phosphorylation of RAB GTPases through hyperactive kinase activity. We found that LRRK2-hyperphosphorylated RABs disrupt the axonal transport of autophagosomes by perturbing the coordinated regulation of cytoplasmic dynein and kinesin motors. In iPSC-derived human neurons, knock-in of the strongly-hyperactive *LRRK2*-p.R1441H mutation caused striking impairments in autophagosome transport, inducing frequent directional reversals and pauses. Knock-out of the opposing Protein Phosphatase 1H (PPM1H) phenocopied the effect of hyperactive LRRK2. Overexpression of ADP-ribosylation factor 6 (ARF6), a GTPase that acts as a switch for selective activation of dynein or kinesin, attenuated transport defects in both p.R1441H knock-in and PPM1H knock-out neurons. Together, these findings support a model where a regulatory imbalance between LRRK2-hyperphosphorylated RABs and ARF6 induces an unproductive “tug-of-war” between dynein and kinesin, disrupting processive autophagosome transport. This disruption may contribute to PD pathogenesis by impairing the essential homeostatic functions of axonal autophagy.

## INTRODUCTION

The leucine-rich repeat kinase 2 (*LRRK2*) gene is a prominent pleomorphic risk locus for Parkinson’s disease (PD), a devastating neurodegenerative disease with debilitating motor and non-motor symptoms. Autosomal dominant missense mutations in *LRRK2* are the most common genetic cause of PD, accounting for ∼5% of familial forms (Healy et al., 2008). In addition, genome-wide association studies have linked noncoding variants in *LRRK2* to risk of sporadic PD.

LRRK2 is a large, multidomain enzyme that includes both a Roc-COR tandem GTPase domain and a catalytic kinase domain. Seven gain-of-function pathogenic mutations either directly (mutations located in the kinase domain) or indirectly (mutations located in the Roc or COR domains) increase LRRK2 kinase activity. The most common pathogenic mutation is *LRRK2*-p.G2019S, located in the kinase domain. Three Roc domain mutations at the p.R1441 hotspot (p.R1441C/G/H) decrease LRRK2 GTPase activity and hyperactivate kinase activity to a similar extent (Steger et al., 2016; Wu et al., 2019). All three mutations induce higher levels of kinase hyperactivity than p.G2019S (Kalogeropulou et al., 2022; Steger et al., 2016). p.R1441 hotspot mutations may also be more highly penetrant than p.G2019S (Haugarvoll et al., 2008; Klein and Westenberger, 2012).

LRRK2 phosphorylates a subset of RAB GTPases (RABs) that coordinate intracellular vesicle trafficking by selectively recruiting effector proteins. Protein Phosphatase 1H (PPM1H) has recently been found to counteract LRRK2 activity by specifically dephosphorylating LRRK2-phosphorylated RAB GTPases (Berndsen et al., 2019). Hyperactive LRRK2 mutations induce increased phosphorylation of RABs, altering their interactions with downstream binding partners. These binding partners include the motor adaptor proteins JNK-interacting protein 3 and 4 (JIP3/4), which selectively interact with LRRK2-phosphorylated RAB proteins and control activation of the molecular motors kinesin and dynein (Bonet-Ponce et al., 2020; Kluss et al., 2022; Waschbüsch et al., 2020). Altered binding to effector proteins induced by LRRK2-mediated phosphorylation of RABs is predicted to alter RAB-dependent intracellular vesicle transport. In recent work, we found that *LRRK2*-p.G2019S decreases the processivity of retrograde axonal AV transport by recruiting JIP4 and inducing inappropriate kinesin activity (Boecker and Holzbaur, 2021; Boecker et al., 2021).

The transport of autophagic vesicles (AVs) in neurons is of particular importance for the maintenance of axonal homeostasis. AVs are formed in the distal axon and at presynaptic sites before undergoing processive retrograde motility toward the soma, driven primarily by the microtubule-associated motor protein cytoplasmic dynein (Cheng et al., 2015; Maday and Holzbaur, 2014; Maday et al., 2012; Neisch et al., 2017; Stavoe et al., 2016). The opposing motor kinesin remains bound to AVs during dynein-driven transport but is auto-inhibited under normal conditions (Boecker et al., 2021; Fu and Holzbaur, 2014; Maday et al., 2012). During retrograde transport, the AV fuses with 1-2 lysosomes, lowering the intra-lumenal pH and activating degradative enzymes (Cason et al., 2022). Importantly, the transport of AVs is tightly-linked to AV maturation and function in cargo degradation (Cason and Holzbaur, 2022; Cason et al., 2021; Wong and Holzbaur, 2014).

Here, we demonstrate that the opposing activities of LRRK2 kinase and PPM1H phosphatase dictate an important regulatory balance for the motor proteins that drive autophagosome transport. We found that neurons expressing the highly-hyperactive *LRRK2*-p.R1441H mutation exhibited a more severe transport deficit than was previously observed in p.G2019S knock-in (KI) neurons (Boecker et al., 2021), suggesting that increased disruptions in autophagosome transport scale with higher levels of LRRK2 kinase activity. In complementary experiments, we found that knock-out (KO) of PPM1H also impaired AV transport, while PPM1H overexpression rescued AV transport in p.G2019S KI neurons, confirming the model that balanced kinase and phosphatase activities are required for normal autophagosome transport. Finally, we found that overexpression of ADP-ribosylation factor 6 (ARF6), which acts as a switch to regulate the association of the activating adaptor JIP3/4 with either dynein/dynactin or kinesin (Montagnac et al., 2009), attenuated defective AV transport in both p.R1441H KI and PPM1H KO neurons. Thus, our data indicate that the balanced interplay among ARF6 activity, JIP3/4 association, and RAB phosphorylation state permits unopposed retrograde autophagosome transport in WT conditions, while hyperactivation of LRRK2 by PD-associated mutations leads to a regulatory imbalance, resulting in an inappropriate “tug-of-war” between dynein and kinesin. Our data robustly link hyperphosphorylation of RABs to defects in neuronal autophagy, an essential homeostatic pathway that has long been implicated in PD pathogenesis.

## RESULTS

### Endogenous LRRK2-p.R1441H disrupts the axonal transport of autophagosomes in iPSC-derived neurons

A point mutation in the kinase domain of LRRK2 (*LRRK2*-p.G2019S) disrupts the axonal transport of autophagic vesicles (AVs) in mammalian neurons. Mutations in the p.R1441 hotspot located in the ROC domain of LRRK2 also cause PD, potentially with higher penetrance (Domingo and Klein, 2018; Haugarvoll et al., 2008; Ruiz-Martínez et al., 2010). Consistent with previous reports, we found that *LRRK2*-p.R1441C causes a higher magnitude of kinase hyperactivity than *LRRK2*-p.G2019S in knock-in (KI) mouse embryonic fibroblasts (MEFs, Figure S1). We measured a ∼2.5-fold increase in RAB phosphorylation over WT for p.R1441C compared to ∼1.5-fold for p.G2019S, representing ∼66% greater hyperactivity of the p.R1441C mutation. As expected, the hyperactive kinase activity induced by either the p.R1441C or p.G2019S mutation was effectively inhibited by the LRRK2 kinase inhibitor MLi-2 (Figure S1).

Given the higher level of hyperactivity induced by a R1441-hotspot mutant, we hypothesized that these mutations would more potently disrupt the axonal transport of AVs. To test this hypothesis, we used human induced pluripotent stem cells (iPSCs) from the KOLF2.1J parental line that were gene-edited to a heterozygous knock-in (KI) of the *LRRK2*-p.R1441H mutation by the iPSC Neurodegenerative Disease Initiative (iNDI) at the NIH (Ramos et al., 2021). We differentiated these iPSCs into excitatory glutamatergic neurons using tetracycline-inducible expression of NGN2 (Pantazis et al., 2022), and live-imaged EGFP-LC3-labeled AV transport in the mid-axon at DIV21 (Figure 1A; Video S1). NGN2-induced neurons (iNeurons) express LRRK2 (Bieri et al., 2019; Boecker et al., 2021; Fonseca-Ornelas et al., 2022) and have been used to study the effect of *LRRK2*-p.G2019S on axonal AV dynamics (Boecker et al., 2021).

**Figure 1.**
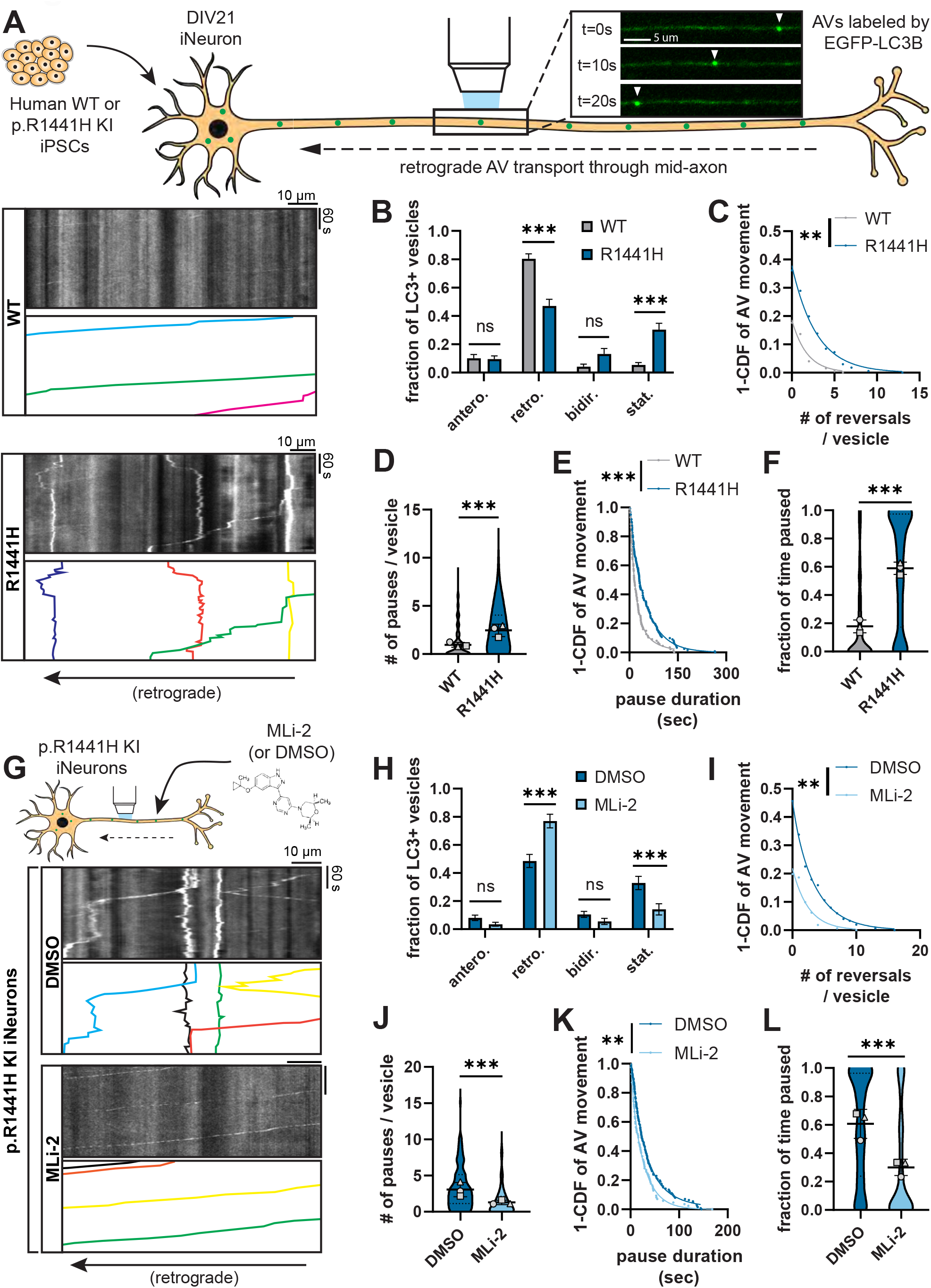
*LRRK2*-p.R1441H knock-in causes kinase-dependent disruption of axonal AV transport. (A) Top, example time lapse images of EGFP-LC3+ vesicles in the mid-axon of a WT iPSC-derived neuron (iNeuron), Left, kymographs of axonal EGFP-LC3+ vesicles in WT and p.R1441H KI iNeurons. Example AV traces are highlighted. (B) Directionality of AVs in WT and p.R1441H KI iNeurons. Antero., anterograde; retro., retrograde; bidir., bi-directional; stat., stationary (mean ± SEM; n = 31 neurons from 3 independent experiments; ns, not significant, p>0.2026; ***p<0.001; two-way ANOVA with Sidak’s multiple comparisons test). (C-E) Directional reversals (C), pause number (D), and pause duration (E) of motile AVs in WT and p.R1441H KI iNeurons (mean ± SD for panel D; n = 168-200 motile AVs from 31 neurons from 3 independent experiments; **p=0.00581; ***p<0.001; mixed effects model analysis, see Methods for specific models used). (F) Fraction of time paused of all AVs in WT and p.R1441H KI iNeurons (mean ± SD; n = 209-246 AVs from 31 neurons from 3 independent experiments; ***p<0.001; mixed effects model analysis). (G) Kymographs of axonal EGFP-LC3+ vesicles in p.R1441H KI iNeurons treated overnight with DMSO or 100 nM MLi-2. (H) Directionality of AVs in p.R1441H iNeurons treated with DMSO or MLi-2 (mean ± SEM; n = 30 neurons from 3 independent experiments; ns, not significant, p>0.7779; ***p<0.001; two-way ANOVA with Sidak’s multiple comparison test). (I-K) Directional reversals (I), pause number (J), and pause duration (K) of motile AVs in p.R1441H KI iNeurons treated with DMSO or MLi-2 (mean ± SD for panel J; n = 153-163 motile AVs from 30 neurons from 3 independent experiments; **p<0.00723; ***p<0.001; mixed effects model analysis). (L) Fraction of time paused of AVs in p.R1441H KI iNeurons treated with DMSO or MLi-2 (mean ± SD; n = 190-221 AVs from 30 neurons from 3 independent experiments; ***p<0.001; mixed effects model analysis). For panels D, F, J, and L, scatter plot points indicate the means of three independent experiments. For panels C, E, I, and K, curve fits were generated using nonlinear regression (two phase decay).

*LRRK2*-p.R1441H caused striking changes in the dynamics of axonal AV transport. When scoring the motility and directionality of AV movement (classifications defined in Figure S2A), we found a marked increase in the stationary fraction of AVs at the expense of the retrograde fraction (Figure 1B). In contrast, we did not observe significant changes in net AV directionality in p.G2019S KI iNeurons in our previous work (Boecker et al., 2021).To further explore which transport parameters might be disrupted by *LRRK2*-p.R1441H, we quantified non-processive motility and pausing behavior of motile AVs in WT and p.R1441H KI iNeurons. p.R1441H KI neurons exhibited reduced AV processivity, with higher number of directional reversals (Figure 1C) and significantly increased Δ run length, a metric of non-processive motility calculated as the difference between the total and net run length of each vesicle (Figure S2C; Δ run length defined in S2B). We also observed a ∼3-fold increase in the number of pauses per vesicle in p.R1441H KI iNeurons (Figure 1D). While this effect was of similar magnitude to that previously observed in p.G2019S KI iNeurons, the average length of each vesicle’s individual pauses (hereafter, pause duration) was increased in p.R1441H KI iNeurons (Figure 1E) but not in p.G2019S KI iNeurons (Boecker et al., 2021).

Thus, expression of the p.R1441H mutation severely disrupted axonal AV transport, with an increased fraction of stationary AVs, an increased number of pauses, and an increase in the pause duration of motile AVs. Combined, these factors manifested as a striking increase in the overall fraction of time that p.R1441H KI AVs spent paused. AVs in p.R1441H KI iNeurons paused for nearly 60% of the time, representing a 3.2-fold increase over WT (Figure 1F). The same quantification performed for AV transport in p.G2019S KI iNeurons (Boecker et al, 2021; data not shown) showed AVs paused for ∼40% of the time (1.97-fold over WT). When normalized to corresponding isogenic WT iNeurons, the p.R1441H effect exceeded that of p.G2019S by ∼61%, appearing to closely correlate with the ∼66% difference in kinase hyperactivity we observed between p.G2019S and a p.R1441 hotspot mutation.

To confirm whether the observed AV transport deficits are LRRK2 kinase activity-dependent, we next tested whether the selective LRRK2 inhibitor MLi-2 (Fell et al., 2015) rescues AV transport in p.R1441H KI iNeurons (Figure 1G). Overnight treatment of iNeurons with 100 nM MLi-2 significantly increased the retrograde fraction of AVs at the expense of the stationary fraction (Figure 1H). Non-processive motility in p.R1441H KI neurons was also rescued, as measured by a reduction of the number of directional reversals and Δ run length (Figure 1I, S2D). Pharmacological LRRK2 inhibition with MLi-2 also reduced pause number and pause duration of motile AVs (Figure 1J-K), rescuing the overall fraction of time spent paused for axonal AVs in p.R1441H KI neurons (Figure 1L). Together, these results are consistent with our previously proposed model of a pathogenic “tug-of-war” between anterograde and retrograde motor proteins induced by hyperactive LRRK2 (Boecker et al., 2021). Importantly, our data also suggest that disruption of AV transport scales in a kinase activity-dependent manner, with greater severity of AV transport defects observed upon expression of p.R1441H as compared to p.G2019S.

### Overexpression of the LRRK2-counteracting phosphatase PPM1H rescues axonal AV transport in p.G2019S KI neurons

Recent work has identified Protein Phosphatase 1H (PPM1H) as the phosphatase counteracting LRRK2 activity through specific dephosphorylation of RAB GTPases (Berndsen et al., 2019). In HEK293 cells transiently expressing LRRK2-p.R1441G, overexpression of GFP-PPM1H^WT^ resulted in reduced levels of phosphothreonine-73 (pT73) RAB10 as compared to overexpression of the catalytically-inactive mutant GFP-PPM1H^H153D^ (Figure S3A). Transient expression of GFP-PPM1H^WT^ in p.G2019S KI MEFs had a similar effect, despite a low transfection efficiency (Figure S3B).

We asked whether enhanced RAB dephosphorylation by PPM1H overexpression could reverse AV transport disruption in neurons expressing hyperactive LRRK2. Using primary cortical neurons from *Lrrk2*-p.G2019S KI mice, we transiently expressed GFP, GFP-PPM1H^WT^, or GFP-PPM1H^H153D^ and visualized AV transport in the mid-axon with mScarlet-LC3. We observed co-transport of GFP-PPM1H with LC3-positive AVs along the axon (Figure 2A, S3C, Video S2). Notably, PPM1H^WT^ overexpression improved retrograde AV processivity in p.G2019S KI neurons compared to either GFP mock or PPM1H^H153D^ transfection (Figure 2B, Video S3). Overexpression of GFP-PPM1H^WT^ significantly decreased the number of pauses, pause duration, and overall fraction of time paused of AVs in p.G2019S KI neurons (Fig. 2C-E). GFP-PPM1H^WT^ overexpression also reduced non-processive AV motility as quantified by the number of directional reversals and Δ run length (Figure 2F-G). We did not observe an effect on AV directionality from PPM1H overexpression (Figure S3D). Together, these data indicate that RAB hyperphosphorylation plays a causal role in the disruption of AV transport by hyperactive LRRK2.

**Figure 2.**
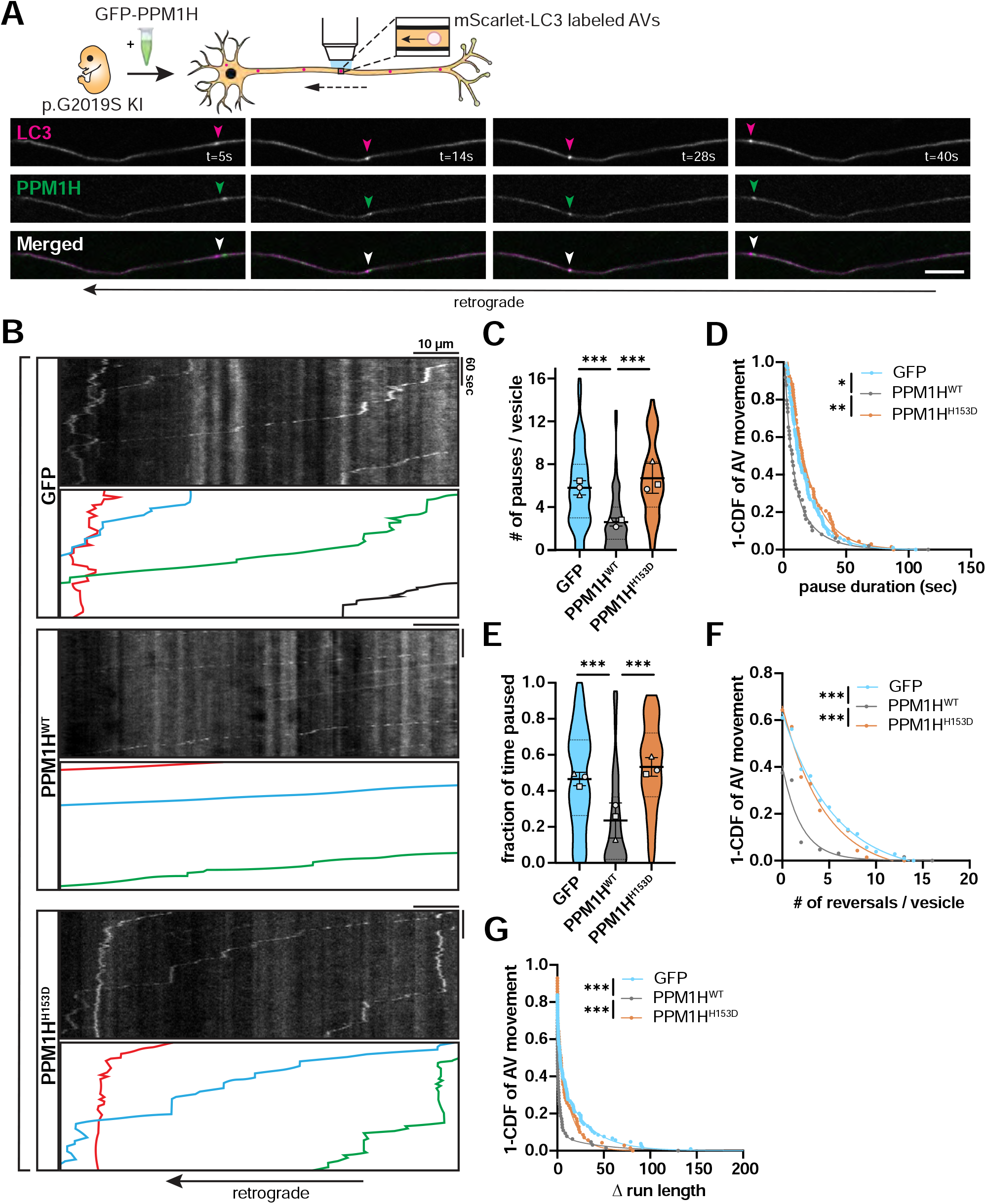
Overexpression of PPM1H rescues AV transport in *Lrrk2*-p.G2019S knock-in mouse cortical neurons. (A) Time lapse images of axonal mScarlet-LC3+ and GFP-PPM1H^WT^+ vesicles in a p.G2019S KI mouse cortical neuron. Scale bar, 10 µm. (B) Kymographs of axonal mScarlet-LC3+ vesicles in p.G2019S KI mouse cortical neurons co-expressing GFP, GFP-PPM1H^WT^, or GFP-PPM1H^H153D^. Example AV traces are highlighted. (C-G) Pause number (C) and pause duration (D) of motile AVs, fraction of time paused (E) of all AVs, directional reversals (F) and Δ run length (G) of motile AVs in G2019S KI mouse cortical neurons transiently expressing GFP, GFP-PPM1H^WT^, or GFP-PPM1H^H153D^ (mean ± SD for panel C and E; n = 66-107 motile AVs (C, D, F, G) and 68-111 total AVs (E) from 21-24 axons from 3 independent experiments; *p=0.01106; **p=0.00144; ***p<0.001; mixed effects model analysis). For panels C and E, scatter plots indicate the means of three independent experiments. For panels D, F, and G curve fits were generated using nonlinear regression (two phase decay for D and F, three phase decay for G).

### PPM1H knock-out phenocopies the effect of hyperactive LRRK2 on AV transport

To confirm the detrimental effect of RAB hyperphosphorylation on axonal AV transport, we next investigated whether PPM1H knock-out (KO) phenocopies the impairment of AV transport by expression of hyperactive LRRK2. Using the KOLF2.1J parental line, we generated PPM1H KO iPSCs through CRISPR/Cas9 gene editing and differentiated them into excitatory glutamatergic iNeurons (Figure S4A). Western blot with a pan-specific antibody for phosphorylated RAB3A, 8A, 10, 35, and 43 showed elevated levels of LRRK2-phosphorylated RAB proteins in PPM1H KO iNeurons (Figure S4B). Similar to expression of the p.R1441H mutation in LRRK2, PPM1H KO dramatically affected AV directionality, with an increase of the stationary fraction at the expense of the retrograde fraction (Figure 3A-B, Video S4). Furthermore, motile AVs in PPM1H KO iNeurons showed a significant increase in the number of pauses as well as a trend toward increased pause duration that did not achieve statistical significance (Figure 3C-D). Together, increased pausing and number of stationary AVs resulted in a marked increase of the overall fraction of time that PPM1H KO AVs spent paused, reaching a similar level as observed in p.R1441H KI iNeurons (Figure 3E). PPM1H KO also phenocopied the effect of hyperactive LRRK2 on non-processive AV motility, increasing the number of directional reversals and Δ run length of motile AVs (Figure 3F-G). To verify whether PPM1H KO and LRRK2 hyperactivity disrupt AV transport through the same pathway, we applied MLi-2 to PPM1H KO iNeurons to inhibit LRRK2 kinase activity (Figure 3H). Compared to DMSO control treatment, MLi-2 rescued AV directionality, with retrograde motility restored to the previously stationary fraction (Figure 3I). MLi-2 furthermore decreased the number of pauses and pause duration of motile AVs, restoring the overall fraction of time paused to levels similar to WT iNeurons (Figure 3J-K, S4C). Non-processive motility of motile AVs, as measured by number of directional reversals and Δ run length was also rescued by LRRK2 inhibition (Figure 3L, S4D). In sum, these results from orthogonal PPM1H model systems provide further evidence that RAB GTPase hyperphosphorylation induces disruption of processive retrograde AV transport.

**Figure 3.**
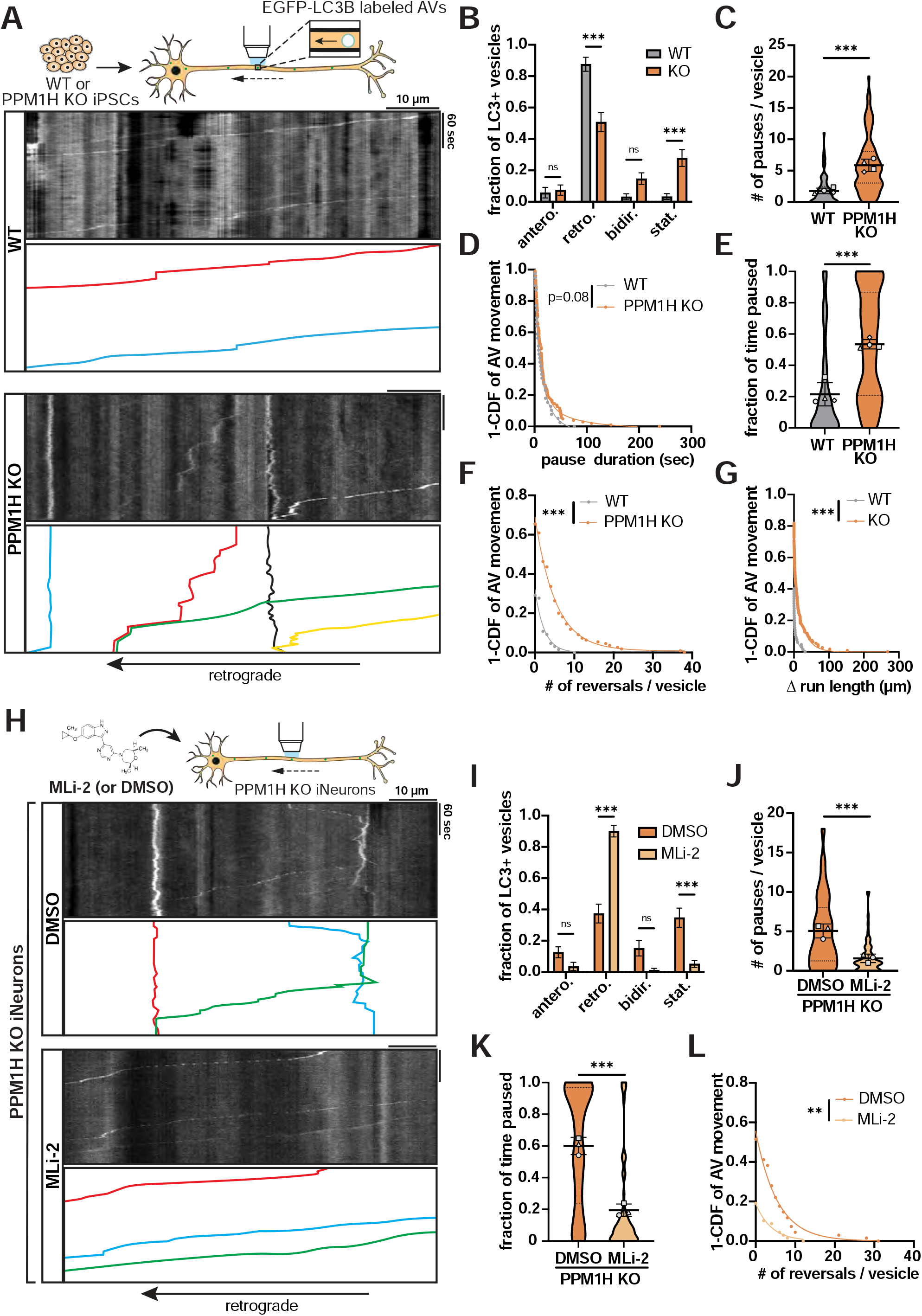
Knock-out of PPM1H causes LRRK2 kinase-dependent disruption of axonal AV transport. (A) Kymographs of axonal EGFP-LC3+ vesicles in WT and PPM1H KO iNeurons. Example AV traces are highlighted. (B) Directionality of AVs in WT and PPM1H KO iNeurons. Antero., anterograde; retro., retrograde; bidir., bi-directional; stat., stationary (mean ± SEM; n = 32-39 neurons from 4 independent experiments; ns, not significant, p>0.1996; ***p<0.001; two-way ANOVA with Sidak’s multiple comparisons test). (C-G) Pause number (C) and pause duration (D) of motile AVs, fraction of time paused (E) of all AVs, and directional reversals (F) and Δ run length (G) of motile AVs in WT and PPM1H KO iNeurons (mean ± SD for panel C and E; n = 65-110 motile AVs (C, D, F, G) and 69-141 total AVs (E) from 32-39 neurons from 4 independent experiments; ***p<0.001; mixed effects model analysis). (H) Kymographs of axonal EGFP-LC3+ vesicles in PPM1H KO iNeurons treated over 72 hours with DMSO or 200 nM MLi-2. Example AV traces are highlighted. (I) Directionality of AVs in PPM1H KO iNeurons treated with DMSO or MLi-2 (mean ± SEM; n = 28-30 neurons from 3 independent experiments; ns, not significant, p>0.0772; ***p<0.001; two-way AMOVA with Sidak’s multiple comparison test). (J-L) Pause number (J) of motile AVs, fraction of time paused (K) of all AVs, and directional reversals (L) of motile AVs in PPM1H KO iNeurons treated with DMSO or MLi-2 (mean ± SD for panel J and K; n = 68 motile AVs (J, L) and 73-104 total AVs (K) from 28-30 neurons from 3 independent experiments; **p=0.00118; ***p<0.001; mixed effects model analysis). For panel C, E, J, and K, scatter plot points indicate the means of three independent experiments. For panels D, F, G, and L, curve fits were generated using nonlinear regression (two phase decay).

### Hyperactive LRRK2 does not disrupt axonal transport of mitochondria

If pathogenic LRRK2 mutations disrupt axonal transport via hyperphosphorylation of RAB proteins, we would predict that the effect would be cargo-specific, such that cargoes whose transport is not regulated by RABs would not be similarly affected. To this end, we tested whether hyperactive LRRK2 affects the axonal transport of mitochondria, a cargo that undergoes both anterograde and retrograde active transport driven by cytoplasmic dynein and kinesin-1 motors but with no known role for RAB GTPases in regulation of their motility (Fenton et al., 2021). Mitochondrial transport in the axon is also of specific interest given the established role of mitochondrial dysfunction in PD pathophysiology (Bose and Beal, 2016).

We visualized mitochondrial transport in the mid-axon of DIV21 p.R1441H KI iNeurons and isogenic WT neurons with Mito-mEmerald (Figure 4A, Video S5). Similar to our previous characterization of DIV21 iNeurons from a different parental iPSC line (Boecker et al., 2020), ∼75% of mitochondria in the mid-axon were stationary or bi-directional in both WT and p.R1441H KI iNeurons. Expression of the p.R1441H mutation neither altered the fraction of motile mitochondria nor perturbed directionality (Figure 4B). We observed a statistically significant but small decrease in axonal mitochondrial density in p.R1441H KI neurons while there was no change in average mitochondrial length (Figure 4C-D), suggesting that the fission/fusion balance of the mitochondrial network was mostly unchanged.

**Figure 4.**
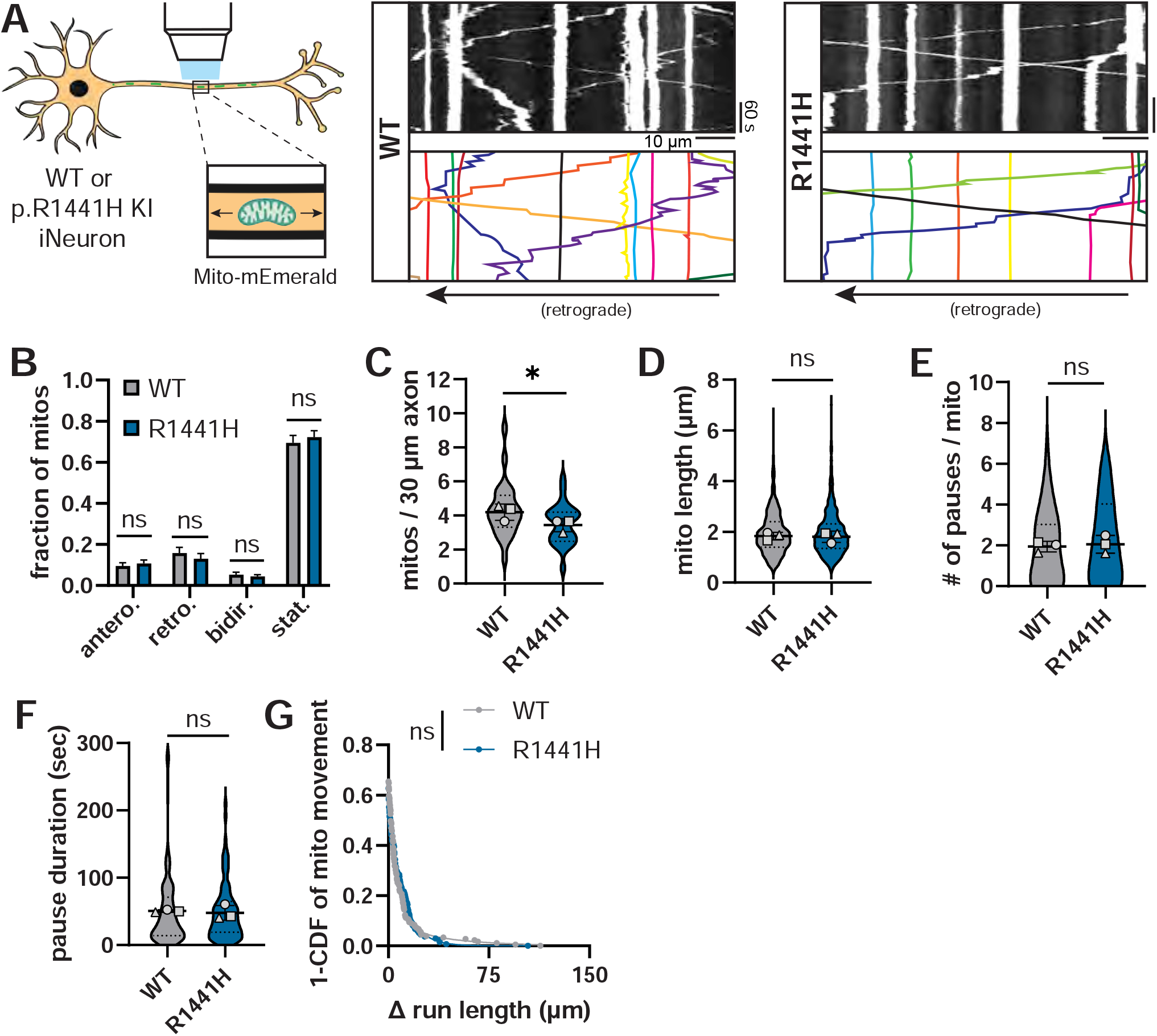
*LRRK2*-p.R1441H does not disrupt axonal mitochondria transport in human iNeurons. (A) Kymographs of axonal mitochondria labeled by Mito-mEmerald in WT and p.R1441H KI iNeurons. Example mitochondrial traces are highlighted. (B) Directionality of mitochondria in WT and p.R1441H KI iNeurons. Antero., anterograde; retro., retrograde; bidir., bi-directional; stat., stationary (mean ± SEM; n = 26 neurons from 3 independent experiments; ns, not significant, p>0.8700; two-way ANOVA with Sidak’s multiple comparisons test). (C-D) Density per 30 µm axon (C) and average length (D) of total population of mitochondria in WT and p.R1441H KI iNeurons (mean ± SD; n = 483-571 mitochondria from 26 neurons from 3 independent experiments; ns, not significant, p=0.2918; *p=0.0348; mixed effects model analysis). (E-G) Pause number (E), pause duration (F), and Δ run length (G) of mitochondria in WT and p.R1441H KI iNeurons (mean ± SD for panels E-F; n = 136-181 motile mitochondria from 26 neurons from 3 independent experiments; ns, not significant, p>0.492; mixed effects model analysis). For panels C-F, scatter plot points indicate the means of three independent experiments. For panel G, a curve fit was generated using nonlinear regression (two phase decay).

To investigate whether LRRK2-p.R1441H disrupts transport of the motile (anterograde, retrograde, and bi-directional) axonal population of mitochondria, we excluded stationary mitochondria from further analysis. We observed no differences in pause number, pause duration, or Δ run length between WT and p.R1441H KI neurons (Figures 4E-G). Furthermore, there were no direction-specific effects on either the anterograde or retrograde population (Figure S5A-D).

We performed similar live imaging of mitochondrial transport in DIV21 *LRRK2*-p.G2019S KI iNeurons and isogenic WT neurons (Figure S6A). In p.G2019S KI neurons, we observed a small but statistically significant decrease in the stationary fraction of mitochondria, but no change in the anterograde, retrograde, or bi-directional fractions (Figure S6B). Axonal mitochondrial density in p.G2019S KI neurons was slightly higher than in WT neurons (Figure S6C), contrasting with the small decrease in mitochondrial density in p.R1441H KI neurons (Figure 4C) and thus suggesting that these changes are unlikely to be a biologically-significant effect of LRRK2 hyperactivity. There was no change in average mitochondrial length (Figure S6D). Similar to our observations with p.R1441H KI neurons, no changes in pause number, pause duration, or Δ run length of the motile population of axonal mitochondria were observed in p.G2019S KI neurons (Figure S6E-G). Likewise, p.G2019S KI neurons did not display direction-specific effects on either the anterograde or retrograde population (Figure S5E-H).

In addition to human iNeurons, we also investigated the effect of hyperactive LRRK2 on mitochondrial transport in primary cortical neurons from *Lrrk2*-p.G2019S KI mice. We used Mito-SNAP to visualize mitochondrial transport in the mid-axon of WT and p.G2019S KI mouse cortical neurons treated with DMSO or 100 nM MLi-2 overnight (Figure S6H). We observed no changes in axonal density, average length, or directionality of mitochondria from either p.G2019S KI or MLi-2 application (Figure S6I-K).

In summary, we found that neither the ROC domain mutation *LRRK2*-p.R1441H nor the kinase domain mutation *LRRK2*-p.G2019S affects axonal mitochondrial transport under basal conditions. Together with our previous observations that *LRRK2*-p.G2019S KI does not impair transport of LAMP1-positive vesicles (Boecker et al., 2021), these results underscore the finding that LRRK2 hyperactivity does not indiscriminately disrupt axonal transport, but rather has a cargo-selective effect on autophagic vesicles.

### Overexpression of the small GTPase ARF6 ameliorates pRAB-mediated disruption of AV transport

Processive retrograde AV transport depends on the tight regulation of the opposing activities of two microtubule-associated motor proteins stably bound to autophagosomes. Both the anterograde motor kinesin-1 and the retrograde motor cytoplasmic dynein (in complex with dynactin) are bound to AVs, but adaptor proteins inhibit kinesin and promote dynein activity in WT neurons (Cason et al., 2021; Cheng et al., 2015; Fu et al., 2014; Maday et al., 2012). In our previous work, we found that hyperactive LRRK2 recruits the motor adaptor JIP4, which specifically binds to LRRK2-phosphorylated RAB proteins, to the AV membrane (Boecker et al., 2021; Bonet-Ponce et al., 2020; Kluss et al., 2022; Waschbüsch et al., 2020). This results in abnormal recruitment and activation of kinesin that disrupts processive retrograde AV transport (Boecker et al., 2021).

The active, GTP-bound form of the small GTPase ARF6 binds to the leucine zipper II domain of JIP4 or its closely related paralog JIP3, promoting interaction of JIP3/4 with dynactin while inhibiting the JIP-kinesin interaction (Montagnac et al., 2009). Our group recently found that overexpression of GTP-locked ARF6^Q67L^ did not affect axonal AV transport in WT rat hippocampal neurons, but expression of GDP-locked ARF6^T27N^ decreased retrograde motility of axonal AVs, supporting that there is a role for ARF6 in the regulation of AV transport (Cason and Holzbaur, manuscript in preparation; available upon request). Notably, while JIP3/4 levels were increased on AVs isolated from mice expressing hyperactive LRRK2, ARF6 levels were not (Boecker et al., 2021). We hypothesized that ARF6 levels become limiting in the presence of hyperphosphorylated RABs, and that overexpression of GTP-ARF6 may restore retrograde AV motility by counterbalancing the abnormally-increased levels of JIP3/4 on the AV membrane (depicted in cartoon in Figure 5A).

**Figure 5.**
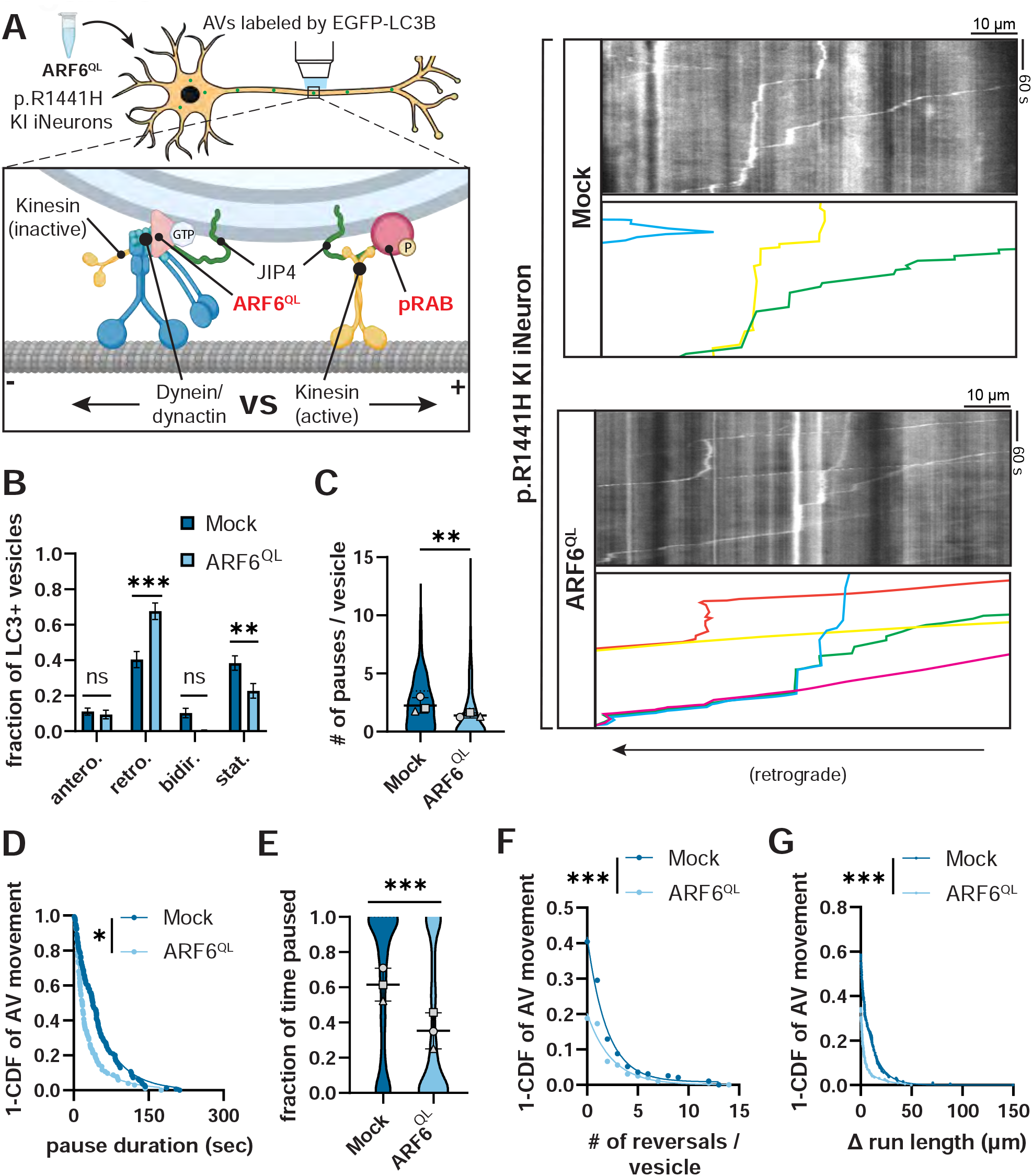
Overexpression of ARF6^Q67L^ ameliorates AV transport deficits in *LRRK2*-p.R1441H KI iNeurons. (A) Kymographs of axonal EGFP-LC3+ vesicles in p.R1441H KI iNeurons transiently expressing CFP (“Mock”) or ARF6^Q67L^-CFP. Example AV traces are highlighted. Cartoon depicts competition between ARF6^Q67L^ and phospho-RAB for binding with JIP4, with opposing directionality of binding partner motors. (B) Directionality of AVs in p.R1441H KI iNeurons transiently expressing CFP or ARF6^Q67L^-CFP. Antero., anterograde; retro., retrograde; bidir., bi-directional; stat., stationary (mean ± SEM; n = 27-28 neurons from 3 independent experiments; ns, not significant, p>0.1492; **p=0.0053; ***p<0.0001; two-way ANOVA with Sidak’s multiple comparisons test). (C-D) Pause number (C) and pause duration (D) of motile AVs in p.R1441H KI iNeurons transiently expressing CFP or ARF6^Q67L^-CFP (mean ± SD for panel C; n = 195-199 motile AVs from 27-28 neurons from 3 independent experiments; *p= 0.0123; **p=0.00261; mixed effects model analysis). (E) Fraction of time paused of all AVs in p.R1441H KI iNeurons transiently expressing CFP or ARF6^Q67L^-CFP (mean ± SD; n = 257-321 AVs from 27-28 neurons from 3 independent experiments; ***p<0.001; mixed effects model analysis). (F-G) Directional reversals (F) and Δ run length (G) of motile AVs in p.R1441H KI iNeurons transiently expressing CFP or ARF6^Q67L^-CFP (n = 195-199 motile AVs from 27-28 neurons from 3 independent experiments; ***p<0.001; mixed effects model analysis). For panels C and E, scatter plot points indicate the means of three independent experiments. For panels D, F, and G, curve fits were generated using nonlinear regression (two phase decay).

To test this model, we transiently expressed the GTP-locked mutant ARF6^Q67L^-CFP in DIV21 LRRK2-p.R1441H KI iNeurons (Figure 5A, Video S6), and visualized AV transport in the mid-axon with EGFP-LC3B. In the CFP mock condition, we again observed dramatic disruption of retrograde AV transport associated with p.R1441H KI. However, overexpression of ARF6^QL^ significantly ameliorated this disruption, increasing the retrograde fraction of AVs at the expense of the stationary fraction (Figure 5B). ARF6^QL^ expression also reduced pausing behavior of motile AVs, reflected by decreases in number of pauses and pause duration (Figure 5C-D). Reduced pausing and stationary fraction resulted in an overall decrease in the fraction of time p.R1441H KI AVs spent paused (Figure 5E). Non-processive motility was also decreased by ARF6^QL^ expression, as quantified by directional reversals and Δ run length (Figure 5F-G). For each metric of AV transport, ARF6^QL^-expressing iNeurons approached but did not reach WT levels (Figure 1), consistent with a partial rescue of the phenotype in p.R1441H KI neurons.

Next, we transiently expressed ARF6^Q67L^-CFP or CFP-only in DIV21 PPM1H KO iNeurons (Figure 6A). Similar to overexpression of ARF6 in p.R1441H KI iNeurons, the retrograde fraction was increased, and the stationary fraction was decreased (Figure 6B). Likewise, we observed significant decreases in number of pauses, overall fraction of time paused, number of directional reversals, and Δ run length. We did not observe a change in the pause duration in PPM1H KO iNeurons as a result of ARF6^QL^ expression (Figure 6C-G). Thus, in two different models of elevated LRRK2-mediated RAB phosphorylation, overexpression of ARF6 partially rescued retrograde axonal transport of AVs. Overall, our findings support a model where increased levels of phosphorylated RABs cause recruitment of additional JIP3/4, which recruits and activates kinesin in the absence of ARF6 regulation. This induces an inappropriately-regulated “tug-of-war” that disrupts processive retrograde AV transport. Overexpression of GTP-ARF6 switches surplus JIP3/4 to an activator of dynein/dynactin instead of kinesin and thereby restores AV processivity.

**Figure 6.**
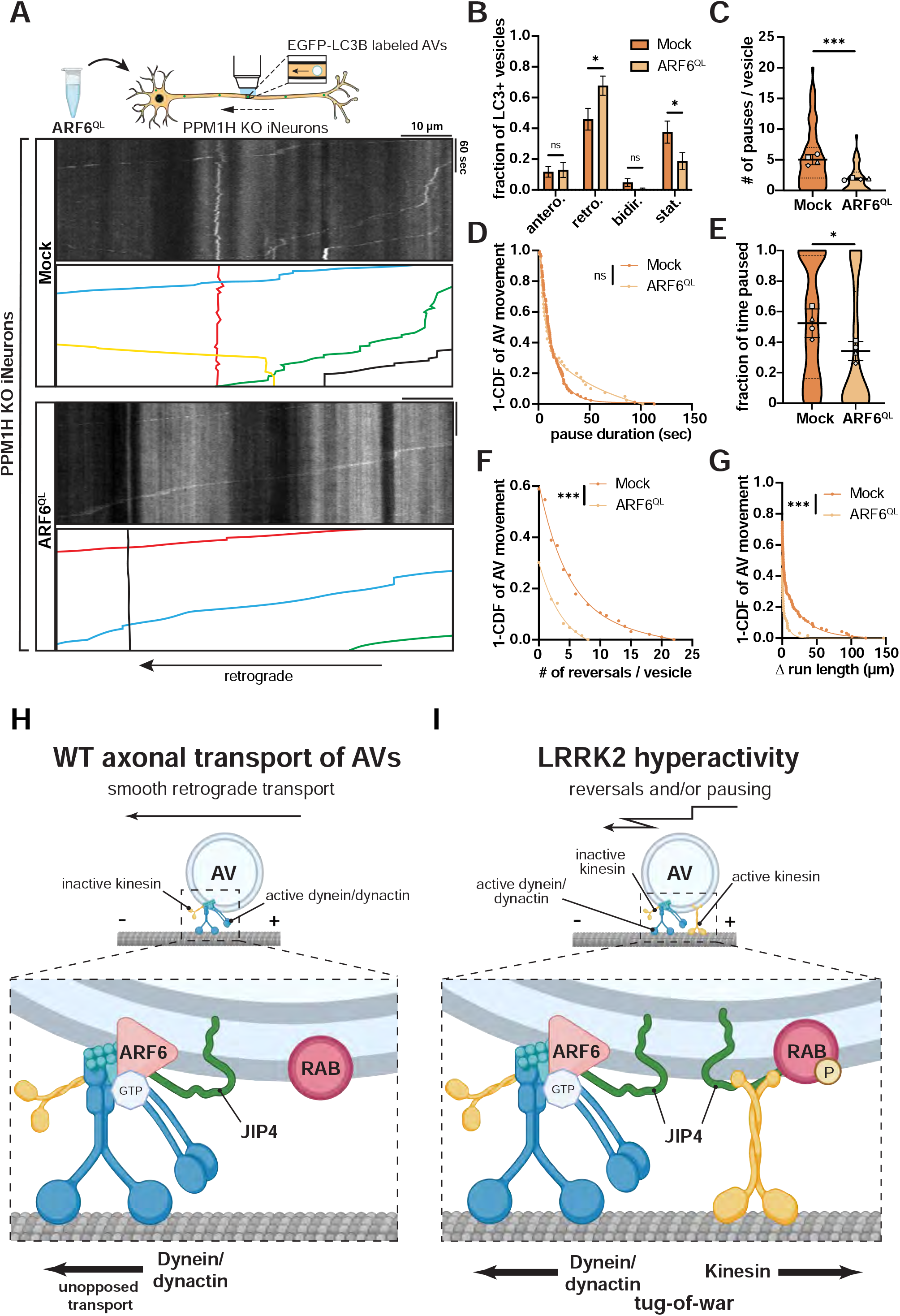
Overexpression of ARF6^Q67L^ ameliorates AV transport deficits in PPM1H KO iNeurons. (A) Kymographs of axonal EGFP-LC3+ vesicles in PPM1H KO iNeurons transiently expressing CFP (“Mock”) or ARF6^Q67L^-CFP. Example AV traces are highlighted. (B) Directionality of AVs in PPM1H KO iNeurons transiently expressing CFP or ARF6^Q67L^-CFP. Antero., anterograde; retro., retrograde; bidir., bidirectional; stat., stationary (mean ± SEM; n = 32-34 neurons from 4 independent experiments; ns, not significant, p>0.9652; *p<0.0410; two-way ANOVA with Sidak’s multiple comparisons test). (C-G) Pause number (C) and pause duration (D) of motile AVs, fraction of time paused (E) of all AVs, and directional reversals (F) and Δ run length (G) of motile AVs in PPM1H KO iNeurons transiently expressing CFP or ARF6^Q67L^-CFP (mean ± SD for panel C and E; n = 63-95 motile AVs (C, D, F, G) and 77-134 total AVs (E) from 32-34 neurons from 4 independent experiments; ns, not significant, p= 0.638; *p= 0.0396; ***p<0.001; mixed effects model analysis). For panels C and E, scatter plot points indicate the means of four independent experiments. For panels D, F, and G, curve fits were generated using nonlinear regression (two phase decay). (H-I) Model: imbalance between ARF6 and pRABs disrupts axonal AV transport in conditions of LRRK2 hyperactivity (H) Under physiologic conditions, ARF6 binds to JIP4, promoting JIP4 binding to dynactin while discouraging the JIP4-kinesin interaction. With active dynein/dynactin and inactive kinesin, AVs undergo smooth retrograde transport in WT neurons. (I) Under pathologic conditions of LRRK2 hyperactivity, increased phosphorylation of RAB GTPases on the AV membrane leads to increased recruitment of JIP4, due to specific binding of JIP4 by RABs in their phosphorylated state. This allows for recruitment and disinhibition of kinesin by JIP4. The presence of active dynein/dynactin and active kinesin on the same organelle results in an unproductive tug-of-war between microtubule-associated motors, leading to non-processive motility and pausing behavior of AVs.

## DISCUSSION

A growing body of evidence links pathogenic LRRK2 mutations to hyperphosphorylation of RAB GTPases, but it remains unclear how phospho-RABs contribute to pathogenesis in PD. Here, we show that increased levels of LRRK2-phosphorylated RAB proteins, resulting from an imbalance in LRRK2 kinase and PPM1H phosphatase activity, disrupt the axonal transport of autophagosomes. Our results further implicate a role for ARF6 in regulating retrograde AV transport (Figure 6H), as expression of GTP-locked ARF6 partially rescued transport deficits. We propose that LRRK2-mediated hyperphosphorylation of RAB GTPases induces dysregulation of the opposing motors dynein and kinesin at the AV membrane in a pathogenic “tug-of-war”. Increased recruitment of JIP3/4 in the absence of enhanced levels of ARF6 inappropriately activates kinesin (Figure 6I), which manifests as increased pausing behavior and non-processive motility of axonal AVs.

We found that *LRRK2*-p.R1441H, a kinase-hyperactivating mutation located in the LRRK2 Roc domain, disrupted axonal AV transport to an even greater extent than the p.G2019S mutation we studied in previous work. While both p.G2019S and p.R1441H increased pause frequency and non-processive motility, only p.R1441H increased average duration of individual pauses and fraction of AVs remaining stationary. Our data suggest that there is an exacerbation of the severity of pausing behavior in the presence of the more hyperactive mutant LRRK2, to the point where a greater fraction of AVs appears to be fully stationary during the confines of a 5-minute live-imaging recording. Combined, the overall fraction of time that AVs spent paused in p.R1441H KI neurons exceeded the effect of p.G2019S by more than 60%. Considering that p.R1441 hotspot mutations cause significantly higher levels of kinase hyperactivity than p.G2019S, our data indicate that the magnitude of AV transport deficits may scale with the level of RAB hyperphosphorylation.

LRRK2 hyperactivity selectively affects axonal cargoes, as autophagosome motility is significantly altered by either hyperactivating mutations in LRRK2 or by loss of the countering phosphatase, but no changes were observed in mitochondrial transport or in lysosome trafficking (Boecker et al., 2021). We hypothesize that AV motility is specifically disrupted because it is regulated by one or more of the RABs that are LRRK2 substrates. In contrast, there is no known role for RAB binding in the mechanism of axonal trafficking of healthy mitochondria, which is controlled by a complex containing the outer mitochondrial membrane protein Miro, the motor adaptor TRAK, and the motor proteins kinesin-1 and dynein/dynactin (Fenton et al., 2021). It is important to note that our live-imaging experiments were all performed at basal conditions, in the absence of pharmacologically-induced mitochondrial damage. It has previously been reported that in the presence of high levels of prolonged mitochondrial stress, p.G2019S KI may alter clearance of axonal mitochondria by impairing degradation of Miro and thereby preventing adequate arrest of motile mitochondria (Hsieh et al., 2016). Further investigation is needed to determine whether such an effect is shared by other pathogenic mutants and whether it can be detected at physiologically-relevant levels of mitochondrial damage.

In PPM1H KO iNeurons, the absence of a key phosphatase opposing LRRK2 activity causes increased levels of pRABs in the setting of LRRK2^WT^ (Figure S4B). In a recent study, loss of PPM1H was found to phenocopy the effects of hyperactive LRRK2 on pRAB-dependent ciliation in mouse cholinergic neurons and astrocytes (Khan et al., 2021). Here, we found that loss of PPM1H caused marked deficits of retrograde AV transport in iNeurons, phenocopying the effect of hyperactive LRRK2 on organelle motility. Defective transport was rescued by treatment with MLi-2, confirming that the alterations in AV transport induced by loss of PPM1H are dependent on LRRK2-mediated RAB phosphorylation. The most likely RABs involved in the LRRK2-dependent impairment of AV transport are RAB10 and/or RAB35, as these RABs are dephosphorylated by PPM1H (Berndsen et al., 2019) and have been implicated in the recruitment of JIP3/4 to organelles (Bonet-Ponce et al., 2020; Waschbüsch et al., 2020). Proteomic and Western blot analyses indicate that RAB10 and RAB35 as well as PPM1H and LRRK2 are all associated with the outer membrane of AVs isolated from brain lysates (Boecker et al., 2021; Goldsmith et al., 2022). However, we cannot exclude the possibility that an as yet unidentified pRab is involved. We note that the AV transport phenotype observed in PPM1H KO neurons more closely resembled the transport deficits seen in p.R1441H KI (this study) rather than the more limited disruption seen previously in p.G2019S KI neurons. This is likely due to the potent increase in levels of pRABs that we and others observed upon PPM1H KO (Khan et al., 2021). Overall, our data imply an important balance between LRRK2 and PPM1H activity in the regulation of AV transport. In both PPM1H KO and p.R1441H KI iNeurons, we found that overexpression of the active, GTP-locked mutant ARF6^QL^ ameliorates defects in AV transport. GTP-ARF6 favors the interaction of JIP3/4 with dynein/dynactin rather than kinesin (Montagnac et al., 2009) and is thus well-positioned as a regulatory switch that promotes uncontested retrograde AV transport. Recent work supports a role for ARF6 in the regulation of AV transport in neurons, as overexpression of the inactive, GDP-locked mutant ARF6^T27N^ impairs retrograde AV transport along the axon (Cason and Holzbaur, manuscript in preparation; available upon request). We propose that hyperphosphorylated RABs disrupt AV transport by recruiting additional JIP3/4 to the AV membrane, leading to erroneous activation of kinesin and causing an unregulated “tug-of-war” between anterograde and retrograde motor proteins, while overexpressed GTP-ARF6 acts as a switch to favor the JIP3/4-dependent activation of dynein/dynactin, thereby restoring highly processive retrograde motility.

How may defects in axonal AV transport be tied to neurodegeneration in PD? The processive transport of AVs is tightly linked to their function as degradative organelles (Cason and Holzbaur, 2022; Cason et al., 2021). Disruption of AV transport induces defective AV acidification and impaired autophagosomal cargo degradation across multiple models of neurodegenerative disease (Boecker et al., 2021; Fu et al., 2014; Wong and Holzbaur, 2014). Recent proteomic work has identified nucleoid-enriched mitochondrial fragments as a main cargo of basal autophagy in neurons (Goldsmith et al., 2022). Thus, defects in neuronal autophagy may lead to the aberrant accumulation of mitochondrial DNA (mtDNA) with the potential to stimulate pro-inflammatory pathways (Riley and Tait, 2020) and thus contribute to the inflammation characteristic of PD progression (Wallings and Tansey, 2019). The degradation of other cargos may also be affected, including the synaptically-enriched protein alpha-synuclein, a key protein in PD pathology and an established substrate of autophagy (Farfel-Becker et al., 2019; Friedman et al., 2012; Vogiatzi et al., 2008). Consistent with this possibility, exacerbated alpha-synuclein aggregation has been linked to expression of hyperactive LRRK2 (Bieri et al., 2019; Ho et al., 2022; MacIsaac et al., 2020; Schapansky et al., 2018; Volpicelli-Daley et al., 2016). Ongoing work to examine the degradation of alpha-synuclein in cellular and animal models of PD should shed further light on this question.

In summary, our data directly link LRRK2-mediated hyperphosphorylation of RAB GTPases to disruption of retrograde AV transport. Mechanistically, we propose that an imbalance between ARF6 and JIP3/4, recruited to the AV membrane by phospho-RABs, disrupts AV transport by inducing an unproductive “tug-of-war” between anterograde and retrograde motor proteins. Our results indicate that the severity of AV transport deficits may scale with the level of LRRK2 hyperactivity, and our data reveal an important balance between LRRK2 kinase activity and the LRRK2-opposing phosphatase PPM1H in regulating AV transport. Together, our work provides new insights on how multiple potential therapeutic targets interact in a pathological mechanism that disrupts axonal autophagy, a homeostatic pathway essential for neuronal health and function.

## ACKNOWLEDGEMENTS

We thank Mariko Tokito for assistance in the cloning of plasmid constructs, Karen Wallace Jahn for assistance with animal models, and Cooper Penner for optimizing our custom KymoTracer analysis software. We thank Michael Ward (National Institutes of Health) and Bill Skarnes (The Jackson Laboratory) for expertise in utilizing resources from the iPSC Neurodegeneration Initiative (iNDI), and the Live-Cell Imaging Facility of the Max Planck Institute for Multidisciplinary Sciences, Goettingen, Germany for technical support. Cartoon schematics were created in part with BioRender.com. Protein structures in graphical abstract were supplied from UniProt.org under the Creative Commons Attribution 4.0 International (CC BY 4.0) License (PDB 7LHT for LRRK2, 7L4J for PPM1H, and 2J5X for ARF6). This work was supported by the National Institutes of Health (1 F31 NS124249-01 and T32-AG-000255 to D.D., R35 GM126950 to E.L.F.H.), the Michael J. Fox Foundation (MJFF-021130 to C.A.B. and E.L.F.H., MJFF-15100 and MJFF-019411 to E.L.F.H.), the German Research Foundation (DFG; BO 5434/2-1 to C.A.B.), and the Howard Hughes Medical Institute (Hanna Gray Fellowship to C.S.E.). The E.L.F.H. laboratory is funded by the joint efforts of The Michael J. Fox Foundation for Parkinson’s Research (MJFF) and the Aligning Science Across Parkinson’s (ASAP) initiative. MJFF administers the grant ASAP-000350 on behalf of ASAP and itself.

## AUTHOR CONTRIBUTIONS

Conceptualization, D.D., C.A.B., and E.L.F.H.; methodology, D.D. and C.A.B.; investigation, D.D., C.A.B., E.M.S., C.S.E.; writing – original draft, D.D., C.A.B., and E.L.F.H.; writing – review and editing, D.D., C.A.B., E.M.S., C.S.E., and E.L.F.H.; funding acquisition, D.D., C.A.B., and E.L.F.H.; supervision, C.A.B. and E.L.F.H.

## STAR METHODS

### KEY RESOURCES TABLE

| REAGENT or RESOURCE | SOURCE | IDENTIFIER |
| --- | --- | --- |
| **Antibodies** |  |  |
| Anti-RAB10, Rabbit Monoclonal | Abcam | Cat# ab237703,<br>RRID:AB_2884879 |
| Anti-RAB10 (phospho T73), Rabbit Monoclonal | Abcam | Cat# ab230261,<br>RRID:AB_2811274 |
| Anti-RAB8A (phospho T72), Rabbit Monoclonal | Abcam | Cat# ab230260,<br>RRID:AB_2814988 |
| Anti-PPM1H, Sheep Polyclonal | MRC PPU reagents and services, University of Dundee, Scotland | #DA018, 3<sup>rd</sup> bleed,<br>RRID:AB_2923281 |
| Anti-Rabbit IgG-IRDye 800CW, Donkey Polyclonal | Li-COR Biosciences | Cat# 926-32213,<br>RRID:AB_621848 |
| Anti-Rabbit IgG-IRDye 680RD, Donkey Polyclonal | Li-COR Biosciences | Cat# 926-68073,<br>RRID:AB_10954442 |
| Anti-Mouse IgG-IRDye 800CW, Donkey Polyclonal | Li-COR Biosciences | Cat# 926-32212,<br>RRID:AB_621847 |
| Anti-Sheep Alexa Fluor 680, Donkey Polyclonal | Thermo Fisher | Cat# A21102,<br>RRID:AB_10374469 |
| **Chemicals, Peptides, and Recombinant Proteins** |  |  |
| MLi-2 | Tocris | Cat# 5756 |
| DMSO | Sigma-Aldrich | Cat# D2650 |
| PLL (mol wt 70,000 – 150,000) | Sigma-Aldrich | Cat# P1274 |
| HBSS (10x) | Thermo Fisher | Cat# 14185-052 |
| 1M HEPES | Thermo Fisher | Cat# 15630-080 |
| 2.5% Trypsin | Thermo Fisher | Cat# 15090-046 |
| Minimum essential medium (MEM) | Thermo Fisher | Cat# 11095-072 |
| Horse serum (heat inactivated) | Thermo Fisher | Cat# 16050-122 |
| Sodium pyruvate | Corning | Cat# 36017004 |
| D-Glucose solution 45% | Sigma-Aldrich | Cat# G8769 |
| GlutaMAX | Thermo Fisher | Cat# 35050061 |
| B27 Supplement | Thermo Fisher | Cat# 17504-044 |
| Neurobasal Medium | Thermo Fisher | Cat# 21103-049 |
| Penicillin-Streptomycin | Thermo Fisher | Cat# 15140-122 |
| AraC | Sigma-Aldrich | Cat# C6645 |
| Lipofectamine 2000 Transfection Reagent | Thermo Fisher | Cat# 11668019 |
| Matrigel Growth Factor Reduced | Corning | Cat# 354230 |
| Essential 8 Medium | Thermo Fisher | Cat# A1517001 |
| mTeSR Medium | Stemcell Technologies | Cat# 85850 |
| Alt-R HiFi Cas9 Nuclease V3 | IDT | Cat# 1081060 |
| ReLeSR | Stemcell Technologies | Cat# 05872 |
| Accutase | Sigma-Aldrich | Cat# A6964 |
| ROCK Inhibitor Y-27632 | Selleckchem | Cat# S1049 |
| Tet System Approved FBS | Takara | Cat# 631107 |
| Knockout Serum Replacement | Thermo Fisher | Cat# 10828010 |
| DMEM/F-12, HEPES | Thermo Fisher | Cat# 11330032 |
| N2 Supplement | Thermo Fisher | Cat# 17502048 |
| Non-essential Amino Acids (NEAA) | Thermo Fisher | Cat# 11140050 |
| Doxycycline | Sigma-Aldrich | Cat# D9891 |
| Poly-L-Ornithine | Sigma-Aldrich | Cat# P3655 |
| BrainPhys Neuronal Medium | Stemcell Technologies | Cat# 05790 |
| Laminin | Corning | Cat# 354232 |
| BDNF | PeptoTech | Cat# 450-02 |
| NT-3 | PeptoTech | Cat# 450-03 |
| Lipofectamine Stem Transfection Reagent | Thermo Fisher | Cat# STEM00003 |
| Microcystin-LR | Sigma-Aldrich | 475815 |
| Halt Protease and Phosphatase Inhibitor Cocktail | Thermo Fisher | 78442 |
| 5-Fluoro-2'-deoxyuridine | Sigma-Aldrich | F0503 |
| Uridine | Sigma-Aldrich | U3003 |
| **Critical Commercial Assays** |  |  |
| BCA Protein Assay Kit | Thermo Fisher | Cat# 23225 |
| Plasmid Maxi Kit | QIAGEN | Cat# 12163 |
| **Experimental Models: Cell Lines** |  |  |
| Human: NGN2 iPSCs | M. Ward (National Institutes of Health)(Wang et al., 2017) | N/A |
| Human: LRRK2-G2019S NGN2 iPSCs (heterozygous) | (Boecker et al., 2021) | N/A |
| Human: LRRK2-WT NGN2 iPSCs | (Boecker et al., 2021) | N/A |
| Mouse: LRRK2-G2019S knock-in MEFs | (Boecker et al., 2021) | N/A |
| Mouse: WT MEFs | MRC PPU reagents and services, University of Dundee, Scotland | N/A |
| Mouse: LRRK2-R1441C knock-in MEFs | MRC PPU reagents and services, University of Dundee, Scotland | N/A |
| Human: KOLF2.1J WT iPSCs | B. Skarnes (Jackson Laboratories, Connecticut) | N/A |
| Human: KOLF2.1J LRRK2-R1441H iPSCs | B. Skarnes (Jackson Laboratories, Connecticut) | N/A |
| Human: KOLF2.1J PPM1H KO iPSCs | This paper | N/A |
| **Experimental Models: Organisms/Strains** |  |  |
| Mouse: LRRK2-G2019S knock-in | Taconic | Model #13940 |
| Mouse: C57BL/6NTac WT | Taconic | Model #B6 |
| **Oligonucleotides** |  |  |
| Gene knockout kit v2 – PPM1H (human) | Synthego | N/A |
| **Recombinant DNA** |  |  |
| Plasmid: PGK EGFP-LC3B | Modified from PGK mCherry-LC3B (gift from Michael Ward, NIH) | Addgene plasmid #193025 |
| Plasmid: PGK ARF6<sup>Q67L</sup>-CFP | This paper | Addgene plasmid #193026 (modified from Addgene plasmid #11387) |
| Plasmid: PGK CFP | This paper | Addgene plasmid #193027 (modified from Addgene plasmid #11387) |
| Plasmid: PGK 4xMito-mEmerald | This paper | Addgene plasmid #193028 (modified from Addgene plasmid #98818) |
| Plasmid: CMV mScarlet-LC3B | Modified from EGFP-LC3B (gift from T. Yoshimori, Osaka University, Japan) | Addgene plasmid #193029 |
| Plasmid: CMV EGFP-PPM1H-WT | MRC PPU reagents and services, University of Dundee, Scotland | #DU62939 |
| Plasmid: CMV EGFP-PPM1H-H153D | This paper | Addgene plasmid #193030 (modified from MRC PPU #DU66215) |
| Plasmid: CMV FLAG-LRRK2-R1441G | MRC PPU reagents and services, University of Dundee, Scotland | #DU13077 |
| Plasmid: PB-TO-hNGN2 | Gift from iPSC Neurodegenerative Disease Initiative (iNDI) & Michael Ward | Addgene plasmid #172115 |
| Plasmid: piggyBac™ transposase vector | Transposagen | N/A |
| **Software and Algorithms** |  |  |
| FIJI (Release 2.9.0) | NIH, USA | https://imagej.net/Fiji |
| Prism 9 | GraphPad | https://www.graphpad.com/scientific-software/prism/ |
| RStudio: Integrated Development for R (2021.09.2 Build 382) | RStudio Team | http://www.rstudio.com/ |
| Matlab R2022a | MathWorks | https://www.mathworks.com/products/matlab.html |
| KymoSuite (custom Matlab script) | (Guedes-Dias et al., 2019) | https://github.com/jnirschl/kinesin-3_guedes-dias_2018 |
| Volocity | PerkinElmer | https://www.perkinelmer.com |
| VisiView 5.0.0.24 | Visitron | https://www.visitron.de/products/visiviewr-software.html |
| Adobe Illustrator 2022 | Adobe | https://www.adobe.com/products/illustrator.html |
| BioRender | BioRender | https://biorender.com/ |
| **Other** |  |  |
| 35 mm #1.5 glass bottom imaging dishes | MatTek | Cat# P35G-1.5-20-C |

## RESOURCE AVAILABILITY

### Lead Contact

Further information and requests for resources and reagents should be directed to and will be fulfilled by the lead contacts, C. Alexander Boecker and Erika L. F. Holzbaur.

### Materials Availability

Unique reagents generated in this study are available from the Lead Contacts with a completed Materials Transfer Agreement.

### Data and Code Availability

Primary data that is presented in this study has been deposited in Zenodo and repository and can be accessed using the Digital Object Identifier 10.5281/zenodo.7178198. The custom MATLAB scripts used in this study to manually track kymographs (KymoSuite) are available at https://github.com/jnirschl/kinesin-3_guedes-dias_2018/tree/master/kymoSuite. Protocols have been deposited to Protocols.io (link will be generated and placed here after formatting of these protocols by Protocols.io staff).

## EXPERIMENTAL MODEL AND SUBJECT DETAILS

### Primary neuron culture

All experiments were performed following protocols approved by the Institutional Animal Care and Use Committee at the University of Pennsylvania. *Lrrk2*-p.G2019S KI mice (model #13940) and B6NTac mice (model #B6) were obtained from Taconic, Cambridge City, Indiana production site. Mouse cortices were dissected from homozygous WT or *Lrrk2*-p.G2019S embryos of either sex at day 15.5. Cortical neurons were isolated by digestion with 0.25% Trypsin and trituration through a small-bore serological pipette. Neurons were plated on 35 mm glass-bottom imaging dishes (P35G-1.5-20-C; MatTek) in Attachment Media (MEM supplemented with 10% horse serum, 33 mM D-glucose and 1 mM sodium pyruvate). After 5 hours, Attachment Media was replaced with Maintenance Media (Neurobasal [GIBCO] supplemented with 2% B-27 [GIBCO], 33 mM D-glucose [Sigma], 2 mM GlutaMAX [GIBCO], 100 U/mL penicillin and 100 mg/mL streptomycin [Sigma]). AraC (1 μM) was added the day after plating to prevent glia cell proliferation. 40% of the media was replaced with fresh Maintenance Media twice per week. Transfections of DIV6 mouse cortical neurons were performed 16-24 hours before imaging using Lipofectamine 2000 Transfection Reagent (ThermoFisher) and 0.9 μg total plasmid DNA.

### i^3^Neuron differentiation

Pre-i^3^Neuron iPSCs (human iPSCs with an integrated doxycycline-inducible mNGN2 transgene in the AAVS1 safe-harbor locus) were a gift from M. Ward (National Institutes of Health, Maryland) and have been described previously (Boecker et al., 2020, 2021; Fernandopulle et al., 2018). Cytogenetic analysis of G-banded metaphases cells showed a normal male karyotype (Cell Line Genetics). Mycoplasma testing was negative. Pre-i^3^N iPSCs were cultured on plates coated with Growth Factor Reduced Matrigel (Corning) and fed daily with Essential 8 media (ThermoFisher). Differentiation of iPSCs into i^3^Neurons was performed using an established protocol (Fernandopulle et al., 2018). In brief, iPSCs were passaged using Accutase (Sigma) and plated on Matrigel-coated dishes in Induction Media (DMEM/F12 supplemented with 1% N2-supplement [GIBCO], 1% NEAA [GIBCO], and 1% GlutaMAX [GIBCO], and containing 2 μg/mL doxycycline. After 72 hours of doxycycline exposure, i^3^Neurons were dissociated with Accutase and cryo-preserved in liquid N2.

### Piggybac-mediated iPSC-derived neuron differentiation

KOLF2.1J-background WT and *LRRK2*-p.R1441H KI iPSCs were a gift from B. Skarnes (Jackson Laboratories, Connecticut) as part of the iPSC Neurodegenerative Disease Initiative (iNDI) and have been described previously (Pantazis et al., 2022). KOLF2.1J-background PPM1H KO iPSCs were generated as described below. Cytogenetic analysis of G-banded metaphases cells showed a normal male karyotype (Cell Line Genetics). Mycoplasma testing was negative. iPSCs were cultured on plates coated with Growth Factor Reduced Matrigel (Corning) and fed daily with Essential 8 media (ThermoFisher). To stably express doxycycline-inducible hNGN2 using a PiggyBac delivery system, iPSCs were transfected with PB-TO-hNGN2 vector (gift from M. Ward, NIH, Maryland) in a 1:2 ratio (transposase:vector) using Lipofectamine Stem (ThermoFisher). After 72 hours, transfected iPSCs were selected for 48 hours with 0.5 μg/mL puromycin (Takara). Induction into neuronal fate with doxycycline and cryopreservation of pre-differentiated neurons was performed as described above (“i^3^Neuron differentiation”).

### Culture and transfection of iPSC-derived neurons

Cryo-preserved, pre-differentiated iNeurons (i^3^Neurons or Piggybac-delivered NGN2 neurons) were thawed and plated on live-imaging dishes (MatTek) coated with poly-L-ornithine at a density of 300,000 neurons per dish. For each experimental condition, cells from at least two different batches of induction were used over three or more independent experimental cultures. iPSC-derived neurons were cultured in BrainPhys Neuronal Media (StemCell) supplemented with 2% B-27 (GIBCO), 10 ng/mL BDNF (PeproTech), 10 ng/mL NT-3 (PeproTech), and 1 μg/mL laminin (Corning). 40% of the media was replaced with fresh media twice per week. For Piggybac-delivered NGN2 neurons, 10 μM 5-Fluoro-2′-deoxyuridine and 10 μM uridine were included at the time of plating to prevent survival of mitotic cells. These drugs were removed 24 hours after plating. Live imaging experiments were performed 21 days after thawing pre-differentiated iPSC-derived neurons (DIV21). On DIV18, iPSC-derived neurons were transfected with Lipofectamine Stem (ThermoFisher) and 1-2.5 μg total plasmid DNA.

## METHOD DETAILS

### Plasmids

Plasmids used include PGK EGFP-LC3 (subcloned from PGK mCherry-LC3B, gift from Michael Ward, National Institutes of Health, Maryland), PGK 4xMito-mEmerald (subcloned from 4xMito-mScarlet-I, Addgene plasmid #98818), PGK ARF6^Q67L^-CFP (subcloned from CMV ARF6^Q67L^-CFP, Addgene plasmid ##11387), PGK CFP (subcloned from CMV ARF6^Q67L^-CFP, Addgene plasmid ##11387), CMV mScarlet-LC3B (subcloned from EGFP-LC3B, gift from T. Yoshimori, Osaka University, Japan, with mScarlet from Addgene plasmid #85054), CMV EGFP-PPM1H-WT (#DU62939, MRC PPU reagents and services, University of Dundee, Scotland), CMV EGFP-PPM1H-H153D (subcloned from pLVX HA PPM1H H153D #DU66215, acquired from MRC PPU reagents and services, University of Dundee, Scotland), CMV FLAG-LRRK2-R1441G (#DU13077, MRC PPU reagents and services, University of Dundee, Scotland), PB-TO-hNGN2 (gift from iPSC Neurodegenerative Disease Initiative (iNDI) & Michael Ward, Addgene plasmid #172115), and piggyBac™ transposase vector (available from Transposagen).

### Generation of PPM1H KO iPSCs

For CRISPR/Cas9 gene editing, KOLF2.1J iPSCs were cultured on Matrigel coated plates in mTeSR medium (StemCell). 800,000 iPSCs were individualized with Accutase and plated onto one well of a Matrigel-coated 6-well plate. iPSCs were then transfected with synthetic sgRNA (Synthego, Gene knockout kit v2 – PPM1H), recombinant Alt-R HiFi Cas9 Nuclease V3 (IDT), and a plasmid encoding for GFP using Lipofectamine Stem Transfection Reagent (ThermoFisher) following the manufacturers’ instructions (12 µL of 1 µM HiFi Cas9 + 12 µL of 1 µM sgRNA in 76 µL OptiMEM combined with 4 µL Lipofectamine Stem in 100 µL OptiMEM). On the following day, iPSCs were split with Accutase and plated in 6 wells of a 6-well plate. 72 hours after transfection, iPSCs were sorted through FACS and 10,000 GFP+ cells were plated on a Matrigel-coated 10 cm dish. Cells were grown for 9 days, then individual colonies were picked. Successful editing was confirmed by Western blot. Karyotype analysis (Cell Line Genetics) demonstrated a normal karyotype of the clone used in this study.

### Live-cell imaging and motility quantification

Primary mouse cortical neurons were imaged on DIV7 in low fluorescence Hibernate E medium (Brain Bits) supplemented with 2% B27 and 2 mM GlutaMAX. iNeurons were imaged on DIV21 in low fluorescence Hibernate A medium (Brain Bits) supplemented with 2% B27, 10 ng/mL BDNF and 10 ng/mL

NT-3. Neurons were imaged in an environmental chamber at 37°C. Recordings of mScarlet-LC3 vesicles in mouse cortical neurons as well as EGFP-LC3 vesicles and mitochondria in WT vs. LRRK2-R1441H iNeurons were acquired on a PerkinElmer UltraView Vox Spinning Disk Confocal system with a Nikon Eclipse Ti inverted microscope, using a Plan Apochromat 60x 1.40 NA oil immersion objective and a Hamamatsu EMCCD C9100-50 camera controlled by Volocity software. Following a scheduled microscope upgrade, the last replicate of the live imaging experiment investigating mitochondrial transport in *LRRK2-*p.R1441H KI iNeurons and all replicates of the experiment investigating the effect of ARF6^Q67L^-CFP overexpression in *LRRK2*-p.R1441H iNeurons were instead performed using a Hamamatsu ORCA-Fusion C14440-20UP camera controlled by VisiView software. Experiments investigating EGFP-LC3 transport in PPM1H KO iNeurons were performed at the Live-Cell Imaging Facility of the Max Planck Institute for Multidisciplinary Sciences, Goettingen, on a Visitron CSU-W1 Spinning Disk Confocal system with a Nikon Ti2 inverted microscope using a Plan Apochromat 60x 1.40 NA oil immersion objective and a Prime BSI sCMOS camera controlled by VisiView software. Axons were identified based on morphological parameters (Boecker et al., 2020; Kaech and Banker, 2006). All time lapse recordings were acquired in the mid-axon (> 300 µm from the soma and > 100 µm from the distal axon terminal) at a frame rate of 1 frame/sec for 5 minutes.

Kymographs of axonal autophagosomes and mitochondria were generated using the Multiple Kymograph plugin (FIJI). Line width was set to 5 pixels. Tracks were traced manually using a custom MATLAB GUI (KymoSuite). Motile autophagosomes and mitochondria were scored as anterograde (net displacement >10 μm in the anterograde direction within the 5-minute time lapse duration), retrograde (net displacement >10 μm in the retrograde direction), or bi-directional (net displacement <10 μm in either direction but total displacement >10 μm). Autophagosomes and mitochondria were scored as stationary if net and total displacement were <10 μm. A pause was defined as a single or consecutive instantaneous velocity value of < 0.083 µm/sec. Bi-directional organelles were included in quantification of pause number, pause duration, directional reversals, and Δ run length, but stationary organelles were excluded. Quantification of the fraction of time paused included all (motile and stationary) autophagosomes. For quantification of Δ run length, the net run length of each vesicle was subtracted from its total run length (Figure S2B). All analyses were performed by a blinded investigator.

Figure legends contain the statistical test used and specific p values for each quantification. For directionality analysis for both AV and mitochondrial transport experiments, two-way ANOVA analysis with Sidak’s test for multiple comparisons was performed using GraphPad Prism 9. For other transport parameters, RStudio version 2021.9.2.382 was used to perform either a linear mixed effects model (LME; R package “nlme”) or a generalized linear mixed model (GLMM; R package “lme4”). The genotype (or, in MLi-2 experiments, the treatment condition) was treated as the fixed effect. The independent experiment/culture and the neuron being recorded from were treated as nested random effects, with the neuron nested within the experiment. Specific models used were chosen based on distribution of each dataset and diagnostic residual plots. In detail, the following models were used: for fraction of time paused, GLMM (binomial family, with the “weights” argument for total time); for pause number, GLMM (Poisson family); for pause duration, GLMM (gamma family, log link function); for reversals, GLMM (Poisson family); for Δ run length, GLMM (gamma family, log link function, with transformation to remove zero values with added constants); for mito density, LME; for mito length, LME. In situations where gamma family GLMMs failed to converge (Figures 1E, 2D, 3D, 4G, 6D, and 6F), adaptive Gauss-Hermite quadrature (nAGQ=0) was used instead of the default Laplace approximation (nAGQ=1). For all quantifications, at least three independent experiments were analyzed.

### Immunoblotting

Neurons or MEFs were washed twice with ice cold PBS and lysed with RIPA buffer (50 mM Tris-HCl, 150 mM NaCl, 0.1% Triton X-100, 0.5%deoxycholate, 0.1% SDS, 2x Halt Protease and Phosphatase inhibitor, 2mg/mL microcystin-LR). Samples were centrifuged for 10 min at 17,000 g, and protein concentration of the supernatant was determined by BCA assay. Proteins were resolved on 8% (LRRK2) - 15% (PPM1H or RAB proteins) acrylamide gels. Proteins were transferred to Immobilon-FL PVDF membranes (Millipore) using a wet blot transfer system. Membranes were then stained for total protein using LI-COR Revert 700 Total Protein Stain. Following imaging of total protein stain, membranes were de-stained and blocked for 5 minutes with Bio-Rad EveryBlot blocking buffer. For PPM1H and pan-specific phosphothreonine RAB blots, membranes were incubated with primary antibody diluted in EveryBlot for 90 minutes at room temperature. For pT73 RAB10 and total RAB10 blots (Figure S3), membranes were incubated with primary antibody diluted in EveryBlot at 4°C overnight. After three washes with TBS (50 mM Tris-HCl [pH 7.4], 274 mM NaCl, 9 mM KCl) with 0.1% Tween-20, membranes were incubated with secondary antibodies diluted in EveryBlot with 0.01% SDS for 1 hr at RT. Following three more washes with TBS with 0.1% Tween-20, membranes were imaged using an Odyssey CLx Infrared Imaging System (LI-COR). Western blots were analyzed with Image Studio Software (LI-COR).

## SUPPLEMENTAL VIDEO TITLES AND LEGENDS

**Video S1. *LRRK2*-p.R1441H knock-in disrupts axonal AV transport in human iNeurons**. Time lapse recording of axonal EGFP-LC3+ vesicles in WT and *LRRK2*-p.R1441H iNeurons. Arrowheads highlight AVs; arrowhead color matches the color of the kymograph traces shown in Figure 1A. Toward the left is retrograde.

**Video S2. Cotransport of GPF-PPM1H and mScarlet-LC3 in *Lrrk2*-p.G2019S knock-in mouse cortical neurons**. Time lapse recording of axonal GFP-PPM1H^WT^**+** and mScarlet-LC3+ vesicles in *Lrrk2*-p.G2019S mouse cortical neurons. Arrowheads highlight axonal AVs. Toward the left is retrograde.

**Video S3. Overexpression of PPM1H**^**WT**^ **rescues AV transport in *Lrrk2*-p.G2019S knock-in mouse cortical neurons**. Time lapse recording of axonal mScarlet-LC3+ vesicles in *Lrrk2*-p.G2019S mouse cortical neurons overexpressing GFP, GFP-PPM1H^WT^, or GFP-PPM1H^H153D^. Arrowheads highlight AVs; arrowhead color matches the color of the kymograph traces shown in Figure 2A. Toward the left is retrograde.

**Video S4. Knock-out of PPM1H causes disruption of axonal AV transport in human iNeurons**. Time lapse recording of axonal EGFP-LC3+ vesicles in WT and PPM1H KO iNeurons. Arrowheads highlight AVs; arrowhead color matches the color of the kymograph traces shown in Figure 3A. Toward the left is retrograde.

**Video S5. *LRRK2*-p.R1441H does not disrupt axonal mitochondria transport in human iNeurons**. Time lapse recording of axonal mitochondria labeled by Mito-mEmerald in WT and p.R1441H KI iNeurons. Arrowheads highlight mitochondria; arrowhead color matches the color of the kymograph traces shown in Figure 4A. Toward the left is retrograde.

**Video S6. Overexpression of ARF6**^**Q67L**^ **ameliorates AV transport deficits in *LRRK2*-p.R1441H KI iNeurons**. Time lapse recording of axonal EGFP-LC3+ vesicles in *LRRK2*-p.R1441H iNeurons overexpressing CFP or ARF6^Q67L^-CFP. Arrowheads highlight AVs; arrowhead color matches the color of the kymograph traces shown in Figure 5A. Toward the left is retrograde.

