## Supplemental Information for "Regulatory imbalance between LRRK2 kinase, PPM1H phosphatase, and ARF6 GTPase disrupts the axonal transport of autophagosomes"

### Figure S1

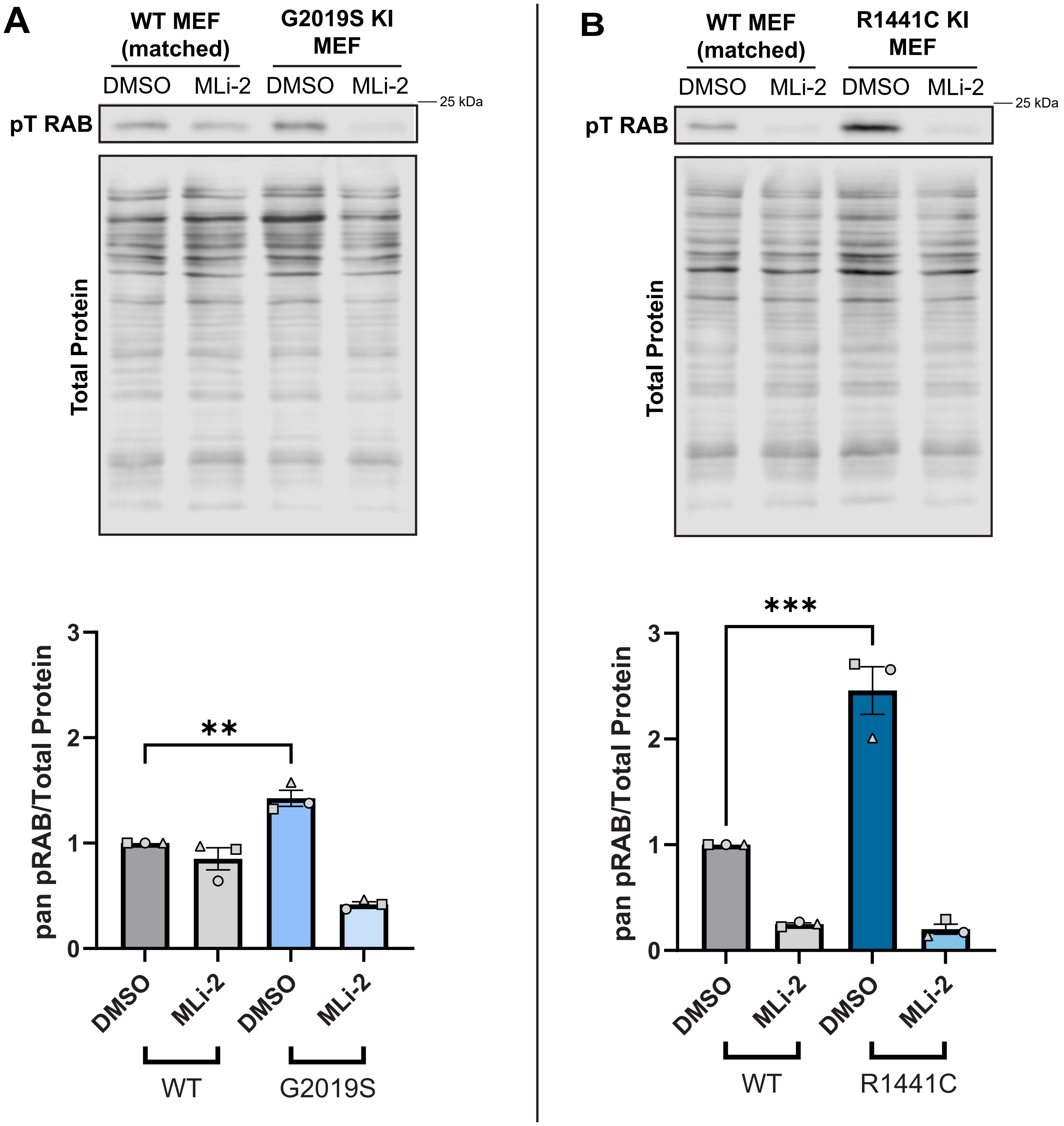

A
WT MEF (matched)
G2019S KI MEF
DMSO
MLi-2
DMSO
MLi-2
25 kDa
pT RAB
Total Protein
pan pRAB/Total Protein
**
0
1
2
3
DMSO
MLi-2
DMSO
MLi-2
WT
G2019S
B
WT MEF (matched)
R1441C KI MEF
DMSO
MLi-2
DMSO
MLi-2
25 kDa
pT RAB
Total Protein
pan pRAB/Total Protein
***
0
1
2
3
DMSO
MLi-2
DMSO
MLi-2
WT
R1441C

**Figure S1. LRRK2 mutant knock-in mouse embryonic fibroblasts (MEFs) have increased levels of LRRK2-phosphorylated RAB proteins. Related to Figure 1.** Representative Western blot and quantification of phosphothreonine RABs in (A) *Lrrk2*-p.G2019S KI MEFs and matched WT MEFs (mean ± SEM; n=3 biological replicates; **p=0.0073; One-way ANOVA with Tukey's multiple comparisons test) and (B) *Lrrk2*-p.R1441C KI MEFs and matched WT MEFs (mean ± SEM; n = 3 biological replicates; ***p=0.0004; One-way ANOVA with Tukey's multiple comparisons test), treated overnight with DMSO or 100 nM MLi-2. Data shown are normalized to total protein and DMSO-treated matched WT condition.

### Figure S2

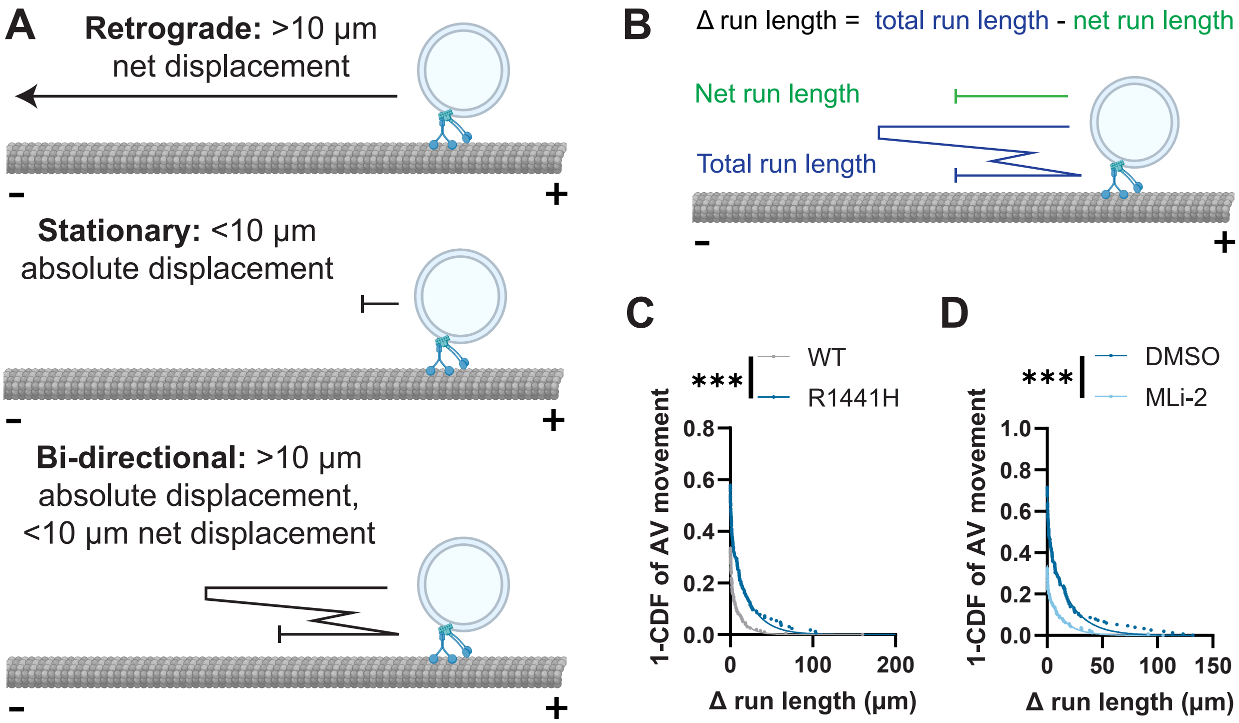
A
Retrograde: >10 µm net displacement
Stationary: <10 µm absolute displacement
Bi-directional: >10 µm absolute displacement, <10 µm net displacement
B
Δ run length = total run length - net run length
Net run length
Total run length
C
WT
R1441H
***
1-CDF of AV movement
Δ run length (µm)
D
DMSO
MLi-2
***
1-CDF of AV movement
Δ run length (µm)

**Figure S2. *LRRK2*-p.R1441H knock-in iNeurons have decreased processivity of AV transport. Related to Figure 1.** (A) Criteria for directionality categorization of AVs in live-imaging experiments. (B) Depiction of Δ run length calculation as a measure of AV transport processivity. (C) Δ run length of AVs in WT and p.R1441H iNeurons (n = 168-200 motile AVs from 31 neurons from 3 independent experiments; ***$p<0.001$; mixed effects model analysis). (D) Δ run length of AVs in p.R1441H KI iNeurons treated with DMSO or MLi-2 (n = 153-163 motile AVs from 30 neurons from 3 independent experiments; ***$p<0.001$; mixed effects model analysis). For panels C and D, curve fits were generated using nonlinear regression (two phase decay).

### Figure S3

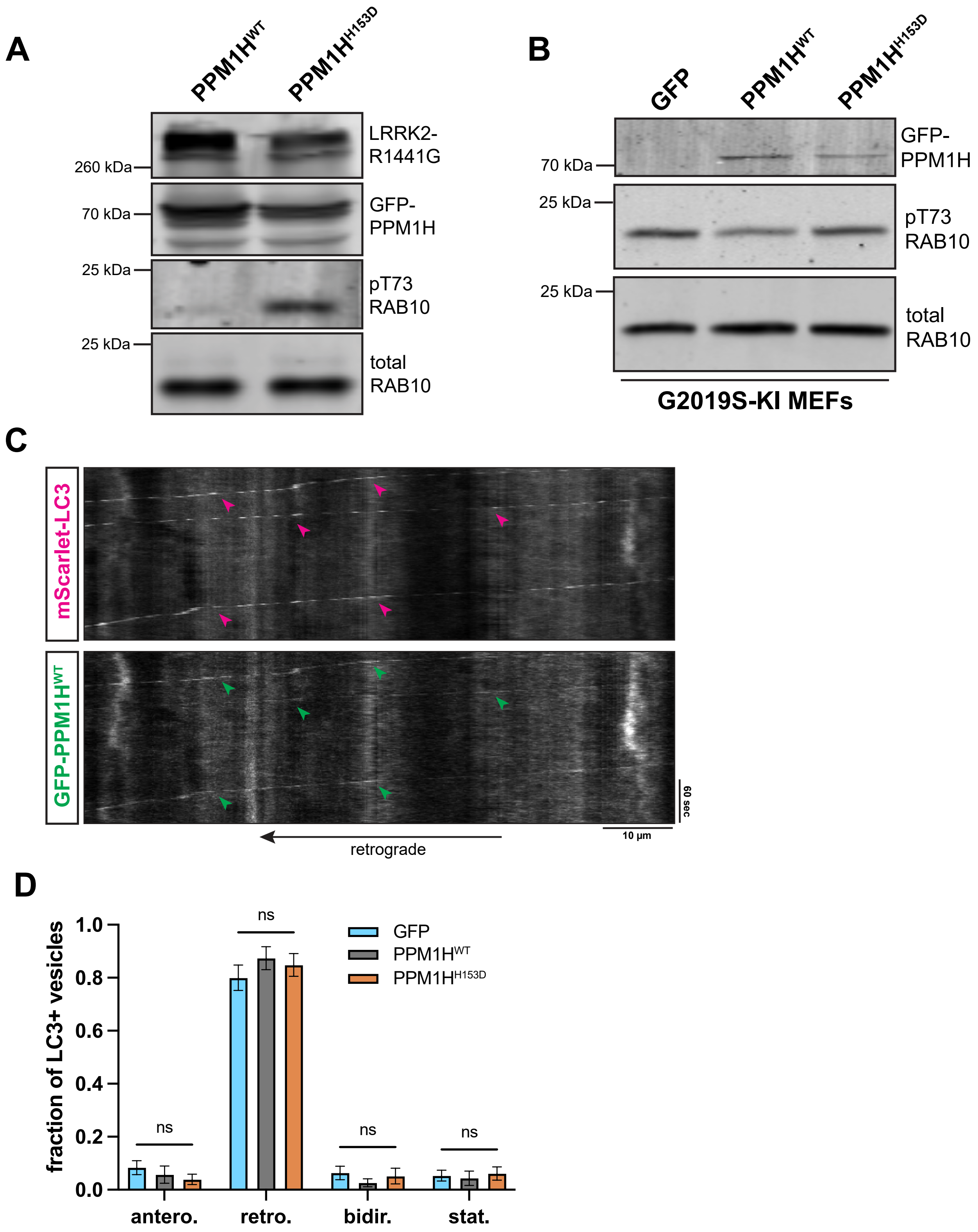
A
PPM1H^WT
PPM1H^H153D
LRRK2-R1441G
260 kDa
GFP-PPM1H
70 kDa
pT73 RAB10
25 kDa
total RAB10
25 kDa
B
GFP
PPM1H^WT
PPM1H^H153D
GFP-PPM1H
70 kDa
pT73 RAB10
25 kDa
total RAB10
25 kDa
G2019S-KI MEFs
C
mScarlet-LC3
GFP-PPM1H^WT
60 sec
10 µm
retrograde
D
fraction of LC3+ vesicles
1.0
0.8
0.6
0.4
0.2
0.0
GFP
PPM1H^WT
PPM1H^H153D
ns
ns
ns
ns
antero.
retro.
bidir.
stat.

**Figure S3. Overexpression of PPM1H rescues AV transport in *LRRK2*-p.G2019S knock-in mouse cortical neurons. Related to Figure 2.** (A) pT73 RAB10 Western blot of HEK 293 cells co-expressing LRRK2-R1441G and GFP-PPM1H$^{WT}$ or catalytically inactive GFP-PPM1H$^{H153D}$. (B) pT73 RAB10 Western blot of *Lrrk2*-p.G2019S knock-in MEFs transiently expressing GFP, GFP-PPM1H$^{WT}$, or GFP-PPM1H$^{H153D}$. (C) Kymographs of axonal mScarlet-LC3+ and GFP-PPM1H$^{WT}$+ vesicles in a mouse p.G2019S KI cortical neuron. Magenta arrowheads point to tracks of mScarlet-LC3+ vesicles, green arrowheads highlight GFP-PPM1H$^{WT}$+ tracks that colocalize with mScarlet-LC3+ tracks. (D) Directionality of AVs in p.G2019S KI mouse cortical neurons transiently expressing GFP, PPM1H$^{WT}$, or GFP-PPM1H$^{H153D}$. Antero., anterograde; retro., retrograde; bidir., bi-directional; stat., stationary (mean ± SEM; n = 21-24 neurons from 3 independent experiments; ns, not significant, p>0.2115; two-way ANOVA with Sidak's multiple comparisons test).

### Figure S4

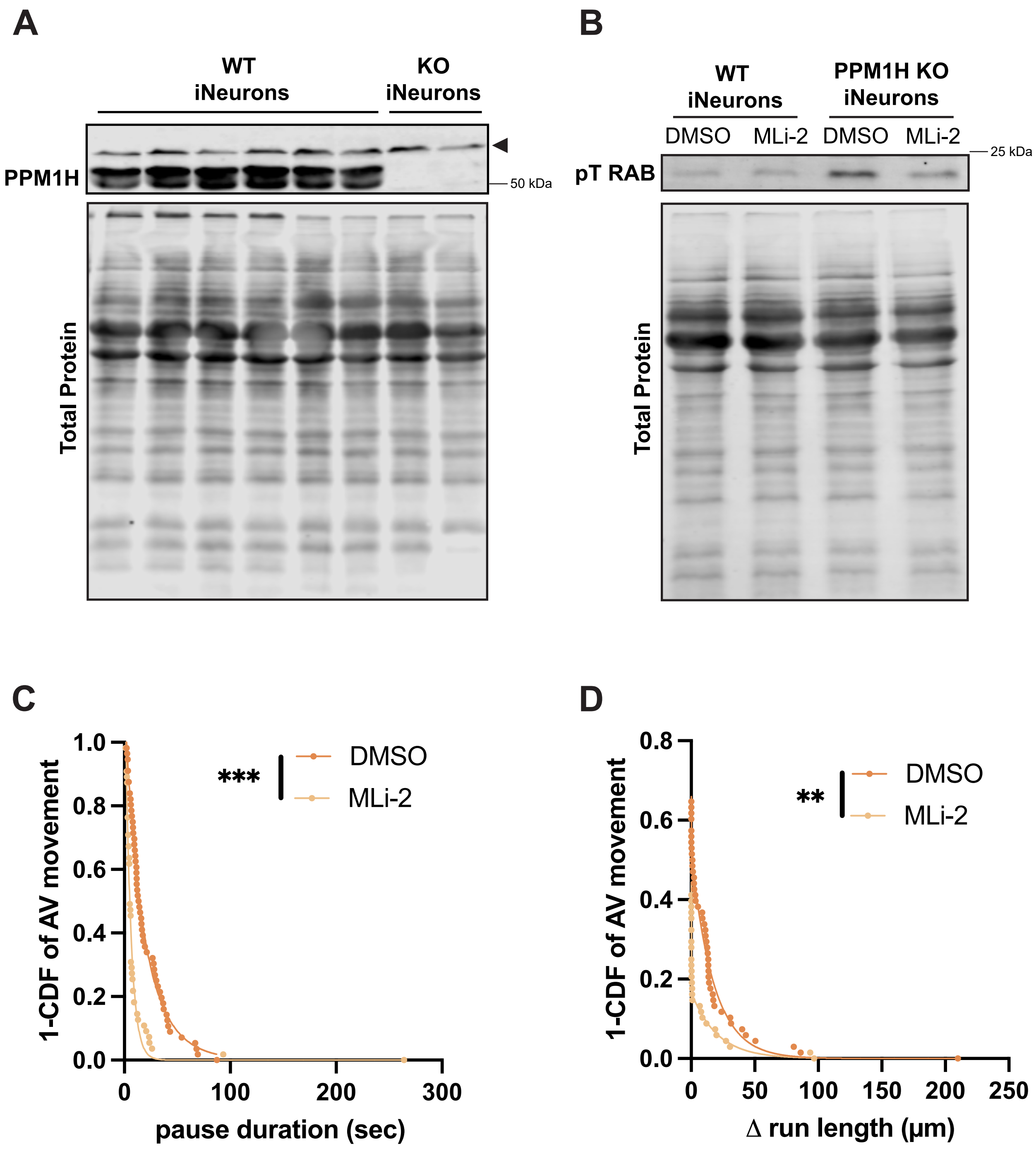
A
WT
iNeurons
KO
iNeurons
PPM1H
50 kDa
Total Protein
B
WT
iNeurons
PPM1H KO
iNeurons
DMSO
MLi-2
DMSO
MLi-2
25 kDa
pT RAB
Total Protein
C
1-CDF of AV movement
***
DMSO
MLi-2
pause duration (sec)
D
1-CDF of AV movement
**
DMSO
MLi-2
Δ run length (μm)

**Figure S4. PPM1H KO iNeurons have increased levels of LRRK2-phosphorylated RAB proteins. Related to Figure 3.** (A) PPM1H Western blot of DIV21 WT or PPM1H KO iNeurons. Arrowhead indicates non-specific band. (B) Pan-specific phosphothreonine RAB Western blot of DIV21 WT or PPM1H KO iNeurons treated with DMSO or 100 nM MLi-2 overnight. (C-D) Pause duration (C) and $\Delta$ run length (D) of motile AVs in PPM1H KO iNeurons treated with DMSO or MLi-2 for 72 hours (n = 68 motile AVs from 28-30 neurons from 3 independent experiments; **p=0.00348; ***p<0.001; mixed effects model analysis). For panels C and D, curve fits were generated using nonlinear regression (two phase decay).

### Figure S5

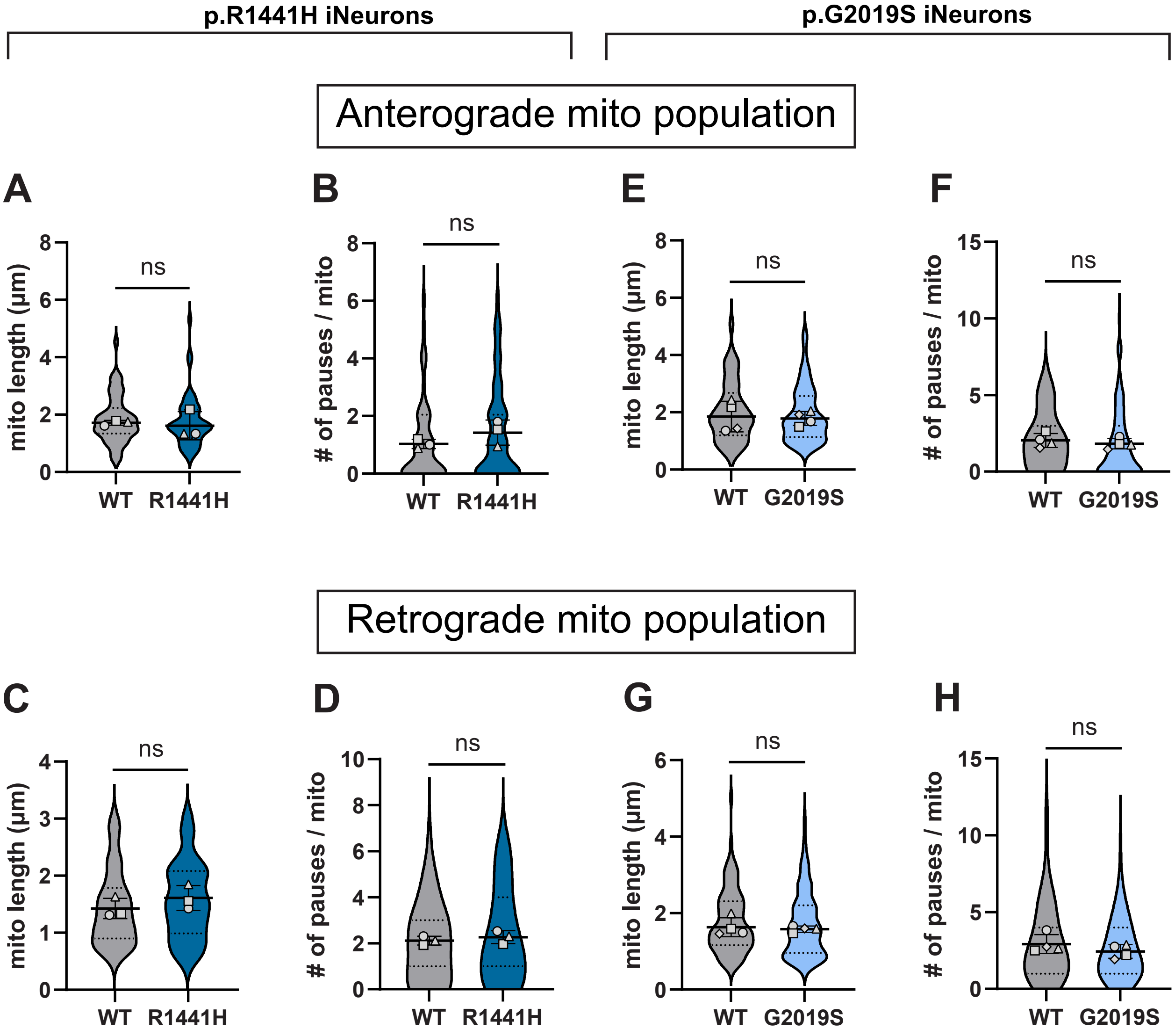

p.R1441H iNeurons
p.G2019S iNeurons
Anterograde mito population
Retrograde mito population
A
B
E
F
C
D
G
H
mito length (µm)
### of pauses / mito
ns
WT
R1441H
G2019S

**Figure S5. Pathogenic *LRRK2* mutations do not disrupt anterograde or retrograde subpopulations of axonal mitochondria. Related to Figure 4.** (A-B) Average length (A) and pause number (B) of anterograde population of mitochondria in WT and p.R1441H KI iNeurons (mean ± SD; n = 51-54 anterograde mitochondria from 26 neurons from 3 independent experiments; ns, not significant, p>0.527; mixed effects model analysis). (C-D) Average length (C) and pause number (D) of retrograde population of mitochondria in WT and p.R1441H KI iNeurons (mean ± SD; n = 63-96 retrograde mitochondria from 26 neurons from 3 independent experiments; ns, not significant, p>0.1233; mixed effects model analysis). (E-F) Average length (E) and pause number (F) of anterograde population of mitochondria in WT and p.G2019S KI iNeurons (mean ± SD; n = 47-91 anterograde mitochondria from 59-60 neurons from 4 independent experiments; ns, not significant, p>0.5598; mixed effects model analysis). (G-H) Average length (G) and pause number (H) of retrograde population of mitochondria in WT and p.G2019S KI iNeurons (mean ± SD; n = 106-162 retrograde mitochondria from 59-60 neurons from 4 independent experiments; ns, not significant, p>0.103; mixed effects model analysis). Scatter plot points indicate the means of 3-4 independent experiments.

### Figure S6

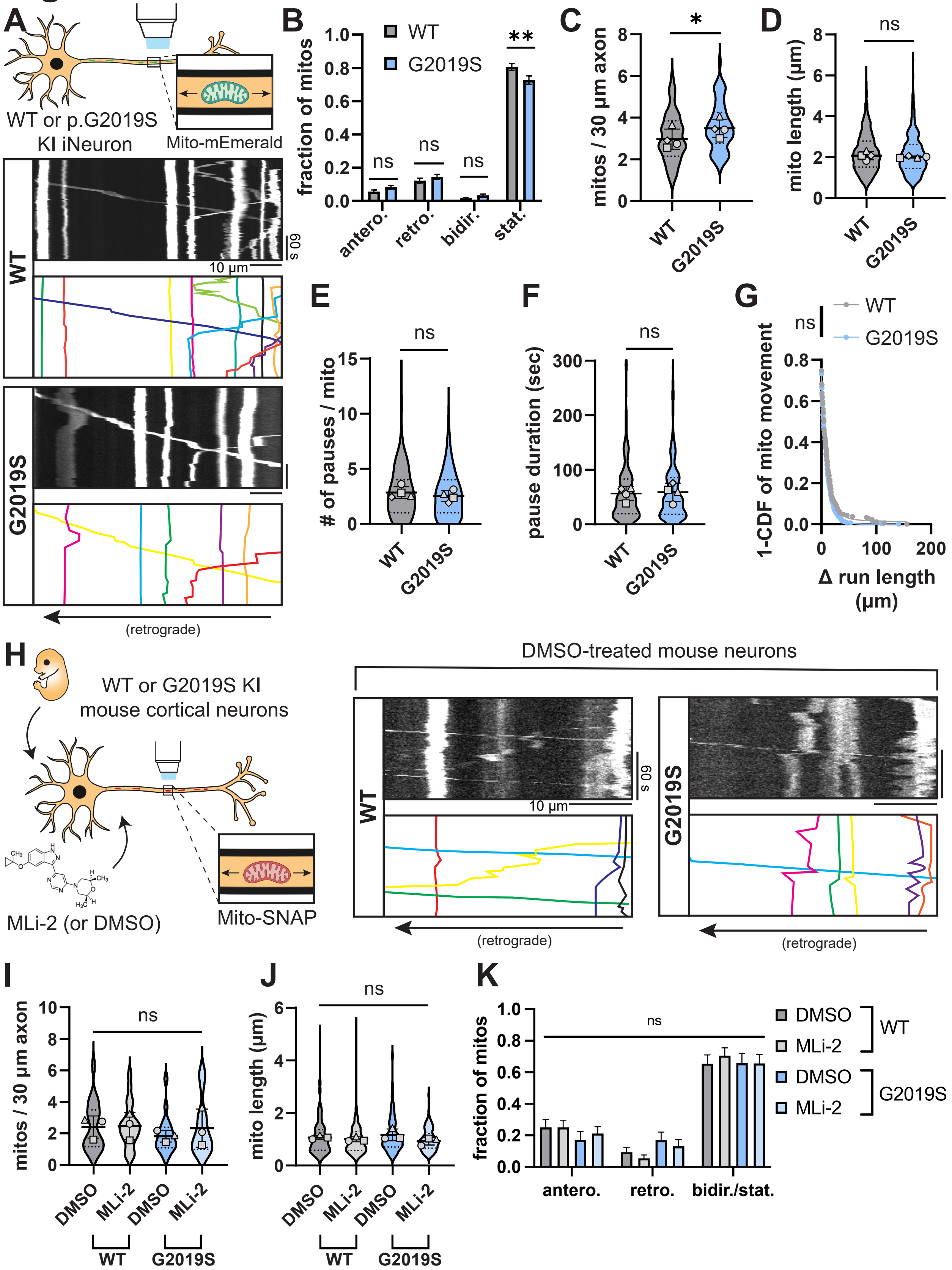
A
WT or p.G2019S KI iNeuron
Mito-mEmerald
WT
G2019S
60 s
10 µm
(retrograde)
B
WT
G2019S
fraction of mitos
antero.
retro.
bidir.
stat.
ns
**
C
mitos / 30 µm axon
*
D
mito length (µm)
E
### of pauses / mito
F
pause duration (sec)
G
1-CDF of mito movement
Δ run length (µm)
H
WT or G2019S KI mouse cortical neurons
MLi-2 (or DMSO)
Mito-SNAP
DMSO-treated mouse neurons
I
mitos / 30 µm axon
DMSO
MLi-2
J
mito length (µm)
K
fraction of mitos
antero.
retro.
bidir./stat.

**Figure S6. LRRK2-p.G2019S does not disrupt axonal mitochondrial transport in human iNeurons or mouse cortical neurons. Related to Figure 4.** (A) Kymographs of axonal mitochondria labeled by Mito-mEmerald in WT and p.G2019S KI iNeurons. Example mitochondrial traces are highlighted. (B) Directionality of mitochondria in WT and p.G2019S KI iNeurons. Antero., anterograde; retro., retrograde; bidir., bi-directional; stat., stationary (mean ± SEM; n = 59-60 neurons from 4 independent experiments; ns, not significant, p>0.6058; **p=0.0012; two-way ANOVA with Sidak's multiple comparisons test). (C-D) Density per 30 μm axon (C) and average length (D) of total population of mitochondria in WT and p.G2019S KI iNeurons (mean ± SD; n = 833-1015 mitochondria from 59-60 neurons from 4 independent experiments; ns, not significant, p>0.1846; *p=0.0105; mixed effects model analysis). (E-G) Pause number (E), pause duration (F), and Δ run length (G) of mitochondria in WT and p.G2019S KI iNeurons (mean ± SD for panels E-F; n = 186-317 motile mitochondria from 59-60 neurons from 4 independent experiments; ns, not significant, p>0.265; mixed effects model analysis). (H) Kymographs of axonal mitochondria labeled by Mito-SNAP in WT and p.G2019S KI mouse cortical neurons treated with DMSO. (I-J) Density per 30 μm axon (I) and average length (J) of mitochondria in WT and p.G2019S KI mouse cortical neurons treated overnight with DMSO or 100 nM MLi-2 (mean ± SD; n = 97-131 mitochondria from 18-21 neurons from 3 independent experiments; ns, not significant, p>0.1924; mixed effects model analysis). Scatter plot points indicate the means of three independent experiments. (K) Directionality of mitochondria in WT and p.G2019S KI mouse cortical neurons treated with DMSO or MLi-2. Antero., anterograde; retro., retrograde; bidir./stat., bi-directional or stationary (mean ± SEM; n = 18-21 neurons from 3 independent experiments; ns, not significant, p>0.4972; two-way ANOVA with Sidak's multiple comparisons test). For panels C-F and I-J, scatter plot points indicate the means of three independent experiments. For panel G, a curve fit was generated using nonlinear regression (two phase decay).
